# Aversive Learning Induces Context-Gated Global Reorganization of Neural Dynamics in *Caenorhabditis elegans*

**DOI:** 10.1101/2025.10.31.685731

**Authors:** Jingting Liang, Sihoon Moon, Sahil Moza, Hyun Jee Lee, Panagiotis E. Eleftheriadis, Juan Chen, Minghai Ge, Maoting Chen, Hang Lu, Yun Zhang

## Abstract

Learning generates experience-dependent changes to the brain. However, how neurons of diverse functions and connectivity reorganize and modulate their activities to generate coherent changes while preserving essential functions is not well understood. Here, we address this question using an aversive olfactory learning paradigm whereby *Caenorhabditis elegans* learns to reduce its olfactory preference for pathogenic bacteria. Using functional imaging during olfactory stimulation in naive and trained animals, we show that, at brain-wide scale, cell type-by-cell type, learning induces context-gated reconfiguration of the organization of neural activity throughout the brain to alter neural responses only during bacteria-discrimination task, while leaving intact bacteria-sensing functions. We found that the context-gated encoding of learning is globally distributed across layers of the nervous system, composed of neurons carrying information of context or learning experience. In particular, aversive training modulates multiple functional connections within the nervous system, including those between sensory neurons and interneurons, as well as those among interneurons, in a context-gated manner. At the systems level, training modulates correlated activity of neural populations; we found that low-dimensional temporal patterns of population activity correlate well with locomotory gaits that express olfactory preferences. Upon training, the rotation and contraction of the low-dimensional neural manifolds shifts the brain into predisposed states for the context-gated display of learned behavior. Because animals encounter unpredictable environments in life, efficient learning about relevant cues while maintaining other functions is essential for survival. Our findings uncover network-level mechanisms that help explain how the brain reorganizes its activity patterns to both encode new experience and preserve essential functions.

## Introduction

Learning engages multiple brain functions, ranging from sensory perception, interneuronal processing, central integration, to motor execution. Consistently, experience-induced modulation of neural activities have been found in various brain regions[1–4]. Previous studies in these brain areas have characterized several network-level strategies for encoding learning. In top-down modulation of functional mapping, signals from the orbitofrontal cortex modulate the activity of the somatosensory cortex when mice learn to respond to changes in the stimulus-reward relationship[5]. In addition, when mice navigate in a simulated space based on the memory of a reward, the choice of locomotory trajectory is encoded by the sequence of activation of the neurons in the parietal cortex, revealing a coding mechanism by activation sequence[6]. A few studies on the lateral amygdala and the dorsal hippocampus have shown that only a portion of the neurons that are capable of encoding memory generate learning-induced changes[2, 7, 8]. This sparse coding strategy allows the population of neurons that share similar connectivity and functions, such as the principal neurons in the dorsal hippocampus, to encode new information with limited interference[2, 7, 8]. Recently, it was shown that previously obtained information that directed a decision-making process in mice was represented by the activity of the neurons in different regions of the brain[9, 10].

While these previous studies identified patterns of brain activity that represent or encode learned information in specific brain regions, we still do not understand whether similar strategies are used to reshape neural activity globally. Moreover, how learning reorganizes neural activity to guide learned behavior while preserving the existing essential functions is largely unknown. These open questions underscore the importance of characterizing neural activity for learning at the brain-wide scale with cell-type resolution in a nervous system with well-defined wiring. Here, we employ the *C. elegans* nervous system as a model to address how learning restructures the activity of diverse neuron classes throughout the brain to generate task-specific modulations in a manner that is compatible with ongoing operations of the nervous system.

## Results

### Multineuronal calcium imaging yields brain-wide neural activities

To study how neural activity encodes learning, we employed an established aversive learning paradigm in *C. elegans*[11, 12]. We previously showed that adult worms grown on their standard laboratory food *Escherichia coli* strain OP50 learned to reduce their innate preference for the smell of the pathogenic bacterium *Pseudomonas aeruginosa* PA14[13] after feeding on PA14 for 4 - 6 hours (Figure S1a)[11, 12]. This form of aversive learning depends on the pathogenesis of the training bacteria and resembles conditioned olfactory or taste aversion that was initially observed in rodents and later widely found in many animals, including humans[14–16]. We quantified the olfactory preference between PA14 and OP50 using an automated behavioral assay that stimulated the worms with alternating odorants of the two bacterial strains and identified turns generated by the worms using machine vision algorithms[11, 17] (Figure S1a). Because attractive odorants suppress the frequency of reorienting turns during chemotaxis to bias movement towards the attractants[11, 17–19], the rates of turns evoked by the bacterial smells were used as an indicator of olfactory preference[11, 17] (Methods and Figure S1).

To characterize brain-wide neural activity for aversive learning of PA14 using *in vivo* calcium imaging (Figure 1a-c), we constructed seven transgenic strains that each stably expressed a genetically-encoded cytoplasmic calcium reporter GCaMP6[20] and a nuclear-localized fluorescent protein mCherry (NLS-mCherry-NLS)[21] in 11 - 21 head neuron classes (Figure S2a, b). To facilitate recording and subsequent analysis, we selected promoters with sparse neuronal expression for each strain. The spatial separation of the labeled neurons in these strains minimizes potential interference among the signals from different cells. The largely stereotyped expression patterns[22] in combination with a collection of transgenes that expressed a blue fluorescent protein (BFP) in specific neuron classes facilitate cell type identification (Figures S2-S3 and Methods). Importantly, these multicellular imaging strains exhibit normal aversive learning ability (Figure S1b - h). We named the neuron populations expressing GCaMP6 and NLS-mCherry-NLS in these transgenic strains based on their cell types[22] as “inter-motor group”, “sensory-inter group I, II, III” and “sensory-inter-motor group I, II, III”, receptively (Figure S2). This strategy enabled us to analyze single-cell resolved neuronal activities in 2,445 recordings from 856 worms that together reported cytoplasmic calcium signals in 78% of 94 classes of *C. elegans* head neurons with high-confidence cell type identification, under both naive and trained conditions, in response to 5 different stimulation patterns (Methods) to characterize how brain-wide neural activities encode learning (Figure 1a - c and Methods).

**Figure 1.**
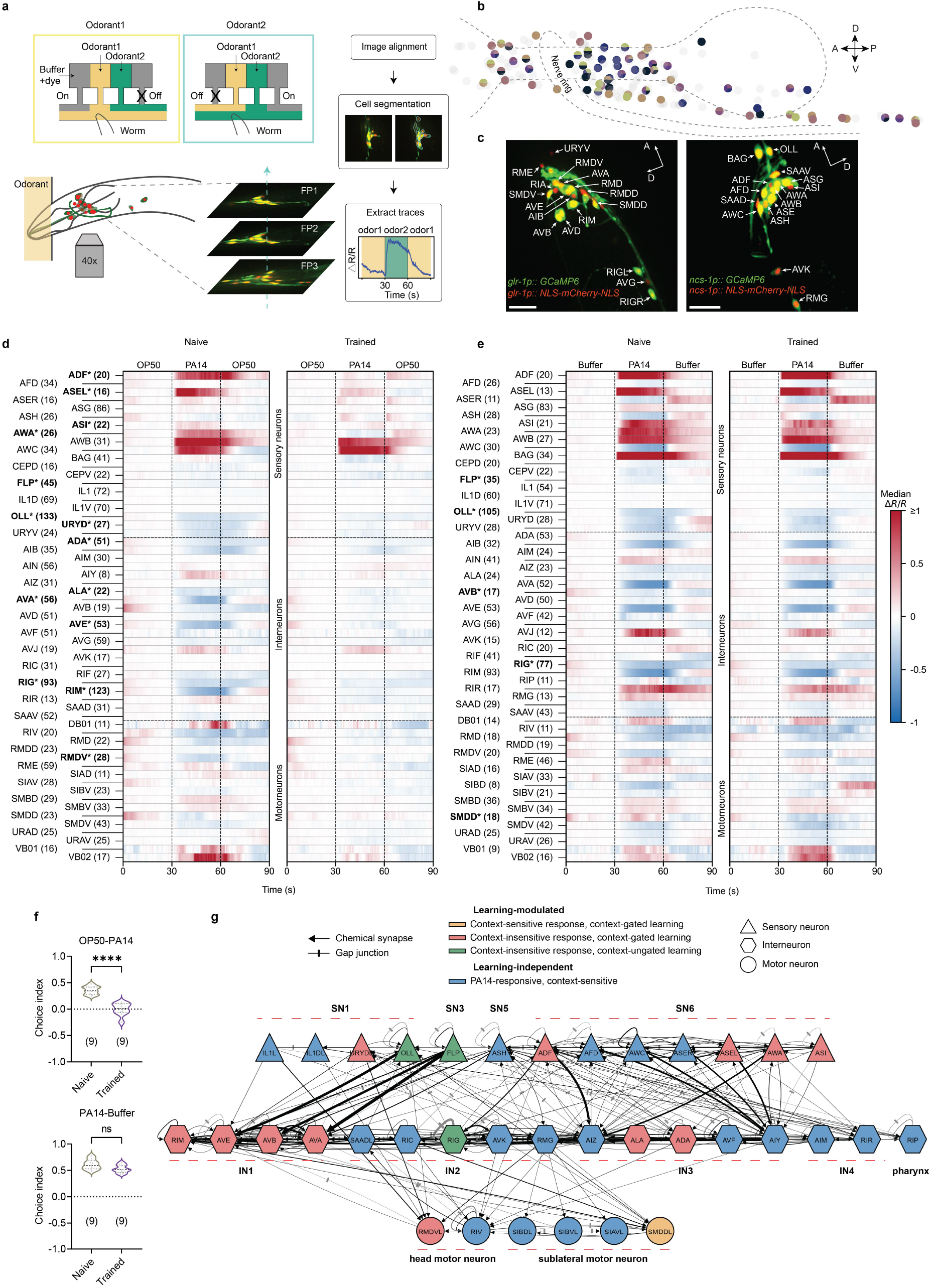
Brain-wide representation of bacterial odorants in *C. elegans* nervous system and context-gated global modulation by aversive training. (a) Schematics showing the procedures for recording and analyzing multineuronal cytoplasmic GCaMP6 signals in defined neuron cell types using a microfluidic device and different stimulation patterns. (b) A schematic showing the positions of all head neurons (gray) and those (highlighted in colors) recorded and analyzed in this study. Dash lines show the positions of the pharyngeal bulb and the nerve ring. Each color highlights one neuron population labeled in one multineuronal imaging strain; one neuron can be labeled in more than one strain. A, anterior; P, posterior; D, dorsal; V, ventral. (c) Sample images for sparse neuronal expression of GCaMP6 and NLS-mCherry-NLS in transgenic imaging strains using the *glr-1* or *ncs-1* promoter. Scale bar, 20 *µ*m. (d, e) Heatmaps for neural activities measured by GCaMP6 signals in identified head neuron classes in response to PA14-conditioned medium in the OP50-PA14-OP50 stimulation pattern (d, discrimination task) or in the Buffer-PA14-Buffer stimulation pattern (e, detection task). Neuron classes recorded in ≥ 8 worms are shown, * and a bold font indicate significantly different neural activities between naive and trained worms in response to either the onset or the removal of PA14-conditioned medium (p *<* 0.05), label-shuffling permutation-test with 5000 iterations followed by Benjamini-Hochberg false discovery rate correction using median of the traces, a number in a parenthesis indicates the smaller number of worms recorded for the neuron class under naive and trained conditions. (f) Violin plots of choice indices between PA14 odorants and OP50 odorants or between PA14 odorants and Buffer tested using droplet assays. Violin plots illustrate the frequency distribution of data points, with the width representing the density of observations. Horizontal lines denote the median (dashed line) and quartiles (dotted lines). The numbers of assays are shown in the parentheses, Student’s *t-*test, asterisks indicate significant difference, **** *p <* 0.0001, ns: not significant. (g) Circuit diagram showing the classification of neurons that encode context, learning, or both, based on the connectome from Cook et al., 2019. For neurons exhibiting strong left–right asymmetry in the connectome, the left side neurons were used for simplicity. ASEL and ASER are shown as two different cell types. Line thickness is linearly proportional to the number of chemical synapses or gap junctions. SN, sensory neuron; IN, interneuron.

### Representation of sensory information is widespread and diverse in *C. elegans* brain

To understand how the experience of feeding on PA14 modulates *C. elegans* brain activity to alter the preference between bacterial odorants, we first characterized the brain-wide neural response to PA14 in naive worms. We strategically chose two different “anchor-odorants”: the OP50-conditioned medium, because OP50 is the default food source that *C. elegans* grows on, and Nematode Growth Medium-buffer that contains little nutritional value. We then presented worms with the PA14-conditioned medium as the “contrast-odorants”. We henceforth refer PA14-conditioned medium, OP50-conditioned medium, and Nematode Growth Medium-buffer as PA14, OP50 and Buffer, respectively. We probed the neural responses with two tasks composed of alternating stimulation of PA14 and OP50 context (OP50-PA14-OP50) or Buffer context (Buffer-PA14-Buffer), referred to as the discrimination task or detection task, respectively (Figure 1).

We found that regardless of the context (OP50 or Buffer), the onset and the removal of PA14 evoked widespread responses with diverse activity patterns (Figure 1d,e, Figure 3a and Figure S4a). First, most sensory neuron classes generated significant responses to PA14 onset or removal under the OP50 context or the Buffer context, including sensory neurons known to detect bacteria-derived chemicals, such as olfactory sensory neurons AWA, AWB, AWC[23] and neurons that had not been fully studied in sensing bacterial cues, such as URY neurons. Most interneuron classes also generated PA14-evoked responses, including interneurons known to respond to sensory stimuli detected by upstream sensory neurons, such as AVA[24], and interneurons with chemosensory functions not fully characterized, such as ADA. Surprisingly, we observed significant PA14-evoked responses in more than 50% of motor neurons under either of these contexts, including head-bending controlling SMDD and SMDV neurons[11, 25, 26]. In addition, some neurons are sensitive to the context, such as ASER, and respond to PA14 differently in OP50 context versus Buffer context (Figure 1d,e and Figure S4a). Second, we found that PA14-evoked neural responses displayed diverse activity patterns with various levels of intensity and temporal dynamics across different layers of the nervous system (Figure 3a). Similarly, stimulating “sensory-inter group I” neurons with OP50 in the context of Buffer (OP50-Buffer-OP50) evoked significant responses in most of the sensory neuron classes with various activity patterns (Figure S5a and Figure S6a). As a control, transitions between Buffer did not elicit significant neural responses (Figure S5b and Figure S6b). Together, our findings are consistent with widespread sensory representations observed in the mammalian brain[10] and demonstrate that the *C. elegans* nervous system is broadly tuned to bacterially generated chemicals. They also provide neuronal basis for processing and integrating sensory information in the layers of sensory, inter- and motor neurons. The diverse patterns of PA14-evoked responses shown here reveal a rich repertoire of neuronal dynamics that is widespread and context-sensitive.

### Brain-wide neural responses are low-dimensional

To characterize network-level activity patterns that emerged from brain-wide neural responses, we applied dimensionality reduction approaches to the time series data. We organized the traces of neuronal signals into a three-dimensional tensor of worms x neurons x trial time (the 30-90 second time window of each trial) (Methods). We applied singular value decomposition (SVD) along the dimension of trial time to generate Temporal Components (TCs). The 3 first TCs together were sufficient to explain about 90% of the total variance (Figure 2a,b) and were thus used to study the neural population dynamics in a reduced-dimension space. These results demonstrate that the widely distributed and diverse neuronal responses to bacterial cues are low-dimensional and share common temporal dynamics.

**Figure 2.**
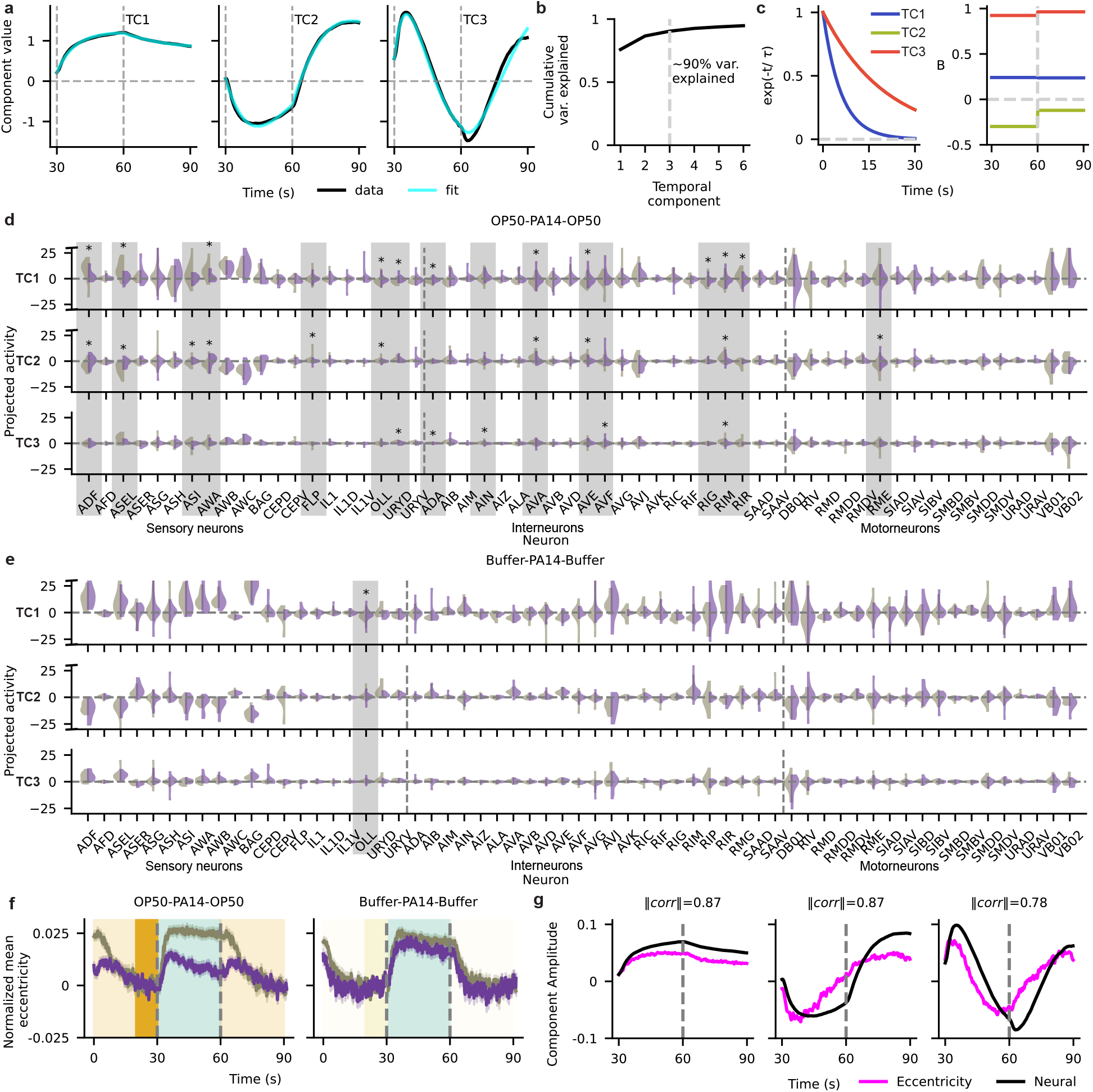
Low-dimensional brain response to PA14. (a) The first 3 Temporal components (TC) extracted from the combined OP50-PA14-OP50 and Buffer-PA14-Buffer datasets (black) overlaid with fits from the linear dynamical system model (cyan, Methods). (b) Nearly 90% of the variance is explained by these 3 components. (c) Decay timescales, *τ* (left), and input strengths, B (right), for TC1, TC2 and TC3. (d,e) Violin-plots showing the distribution (kernel density estimates) of the neural activity projected to TC1, TC2 and TC3 for naive (gray) and trained (purple) conditions for OP50-PA14-OP50 stimulation pattern (d, discrimination task) and for Buffer-PA14-Buffer stimulation pattern (e, detection task). Neurons that are significantly different for a specific TC in naive and trained conditions are marked by an asterisk and a gray shade. Angled dash on the axis spine indicates a truncated axis. (f) Average eccentricity values normalized to the last 10 seconds of the context-odorants (OP50 or Buffer for OP50-PA14-OP50 and Buffer-PA14-Buffer stimulation patterns, respectively), identical to the normalization used for the neural data. Shade: SEM. (g) Behavioral components (BC1, BC2, BC3) calculated from eccentricity values in magenta and temporal components (from a) in black showing a high absolute correlation.

We observed that the shapes of TC1 and TC2 were clearly aligned to the stimulus transition points, suggesting that these components were strongly input-driven and that they illustrated either the direct sensory responses or the downstream activities evoked by upstream sensory responses or the integration of both signals. To capture their properties quantitatively, we modeled the temporal components as a linear dynamical system *Ax* + *Bu* (Methods), where *u* is the step-function odorant input, *B* is input-scaling, with which the odorant input transforms into neural activity and *A* is an interaction term between the components to capture the internal dynamics of the network. We saw that input-scaling had different values for two classes of odorants for these three components (Figure 2a-c). TC1 and TC3 had similar values of input-scaling between the contrast odorants (PA14) and the anchor odorants (OP50 and Buffer) and TC2 had strong asymmetric input scaling between these two odorant classes. Together, these shapes encode ways to both detect (TC1 and 3) and differentiate (TC2) between odorant stimulations on the single trial level. In addition, we found that TC1 and TC2 had the same timescale, while TC3 had a longer timescale (Figure 2c and Methods), pointing to the maintenance of stimulation information at different timescales and across stimulation periods. By comparing the magnitudes of *A* and *B* for the different components individually (Figure 2c), we deduced that the dynamics of TC1 can be explained as being strongly input-dependent, and TC2 and TC3 can be explained as a mixture of direct inputs and cross-interactions with other components.

To further understand the information represented by the temporal components of the neuronal populations, we examined neurons that displayed high scores in naive worms (Figure 2d,e). The sensory neurons with high loadings of TC1 include those that detect metabolic products from different bacteria, including PA14[11, 27, 28]. Several interneurons that receive direct synaptic input from the sensory layer (*e*.*g*. AIB)[29], as well as some downstream motor neurons (*e*.*g*. SMDD/V that control head bending[26, 30, 31]) also show high loadings of TC1 (Figure 2d,e). These results suggest that sensory-evoked neural signals travel deeply into the downstream layers of the neural network, including motor neurons to potentially tune their locomotion-controlling activities. Most neurons exhibiting high loadings of TC2 also strongly contribute to TC1, suggesting that TC2 also represents primary sensory responses, consistent with our analysis of the shapes above. Several neurons showed small loadings of TC3, which weakly contributed to the total variance explained (Figure 2b,d,e).

### Low-dimensional population activity encodes locomotory gait expressing behavioral preference

Next, we sought to address how the brain-wide neural responses to the bacteria discrimination task and detection task represent behavior. Worms swim by continuously bending their bodies and a higher rate of turns, which lead to reorientation, indicates a lower preference to the presented stimulus[11, 17, 25]. Thus, we quantified locomotory gait of the swimming worms in the olfactory droplet assay using the degree of bending, measured by the eccentricity of an ellipse fitted to the worm body. Small eccentricity values were used as a proxy for strong body bends and turns (Figure S7 and Methods). We found that switching to PA14 from either OP50 or Buffer suppressed turns in naive worms and that training with PA14 released the suppression, but only in the context of OP50 in the discrimination task, not in the context of Buffer in the detection task (Figure 1f). These training-dependent task-specific effects were fully captured by quantification of eccentricity values (Figure 2f). Thus, we used eccentricity to quantify locomotory gait that reveals behavioral preference. We performed principal component analysis (Methods) on the eccentricity values in naive and trained worms. We found that the first three components explained 44% of the total variance. Strikingly, the first three TCs for the population neural response are highly correlated with the first three components of eccentricity (Figure 2g), consistent with encoding of the behavioral preference by the network neural responses evoked by PA14. Brain-wide neural encoding of locomotion has been demonstrated in both vertebrate and invertebrate brains[10, 32–34]. Our findings on the encoding of behavioral preferences by the population neural activity and global representation of sensory responses support widely distributed sensory-motor integration in the nervous system that subserves experience-dependent modulation for learning.

### Aversive training globally modulates brain activity in a context-gated manner

To characterize how aversive training with the pathogenic bacterium PA14 reorganizes the brain activity in *C. elegans*, we compared brain-wide calcium imaging signals from naive and trained worms (Figure 1d,e and Methods). We found that aversive training altered the neural responses to PA14 in the discrimination task with OP50 context in neurons across layers of the nervous system. Specifically, 35% of sensory neuron types, 27% of interneuron types and 6% of motor neuron types exhibited significant learning-dependent changes in the amplitudes of their GCaMP6 signals evoked by the onset or the removal of PA14 in the discrimination task (Figure 1d,e). The critical role in regulating aversive learning of PA14 has been demonstrated for 29% (ADF, AWA, AVA, RIM[12, 28, 35], 4/14) of the neuron cell types showing learning-dependent modulation, supporting the importance of the identified neurons in the aversive learning and revealing global encoding of the learning in the worm brain. In contrast, in the detection task with Buffer context only 5 cell types exhibited significant changes in their calcium signals with only 3 of these cell types displaying similar learning-dependent changes under both contexts (Figure 1d,e). As a control, stimulation of Buffer to Buffer transitions did not evoke different neural responses in naive and trained worms (Figure S5b and Figure S6b), consistent with the behavioral outputs (Figure S5d). These results demonstrate that the aversive training-induced modulation of neural activity is systematically displayed in a context-gated manner, which enables task-specific neural responses that strongly reduce the preference for PA14 over OP50 without altering the preference of PA14, a food, over Buffer, a non-food stimulus. Thus, our results reveal that the context-gated modulation of neural response allows the encoding of new information while preserving the existing function of the brain.

We classified neurons based on whether they respond to PA14 in naive worms, whether their naive responses to PA14 differ in the discrimination task and detection task, whether their PA14 responses are modulated by learning, and whether the learning-induced modulation is context-gated (Figure S5h and Methods). This analysis identified four categories of neurons encoding context, learning, or both (Figure 1g). We found that neurons in different classification categories are distributed across different layers of the nervous system, including sensory neurons, interneurons, and motor neurons. In addition, neurons in different categories interconnect with each other. These analysis suggests integrated encoding of context and learning by neurons in different layers of the nervous system with different connectivities.

Next, we examined how aversive training with PA14 altered the patterns of population neural activity by comparing the loadings of the temporal components (TCs) between naive and trained worms in the bacteria discrimination task and detection task. We observed widespread changes to sensory neurons and interneurons in both TC1 and TC2. First, we found that 85% of the neurons that showed significant learning-dependent changes in the amplitudes of their calcium signals in the discrimination task also significantly altered their loadings of at least one TC (Figure 2d,e), demonstrating that the modulation of these neurons strongly contributes to the reorganization of the network activities and that the learning-induced reorganization of population activities occurs largely in the same low-dimensional space. Compared with 16 cell types that showed learning-induced changes in their TC loadings in the discrimination task, only 1 cell types, OLL, changed the TC loadings in the odorant detection task and displayed similar changes in both contexts (Figure 2d,e). Largest changes in the odor discrimination task corresponded to amphid neurons group (ADF, AWA, ASEL, ASI) and layer 1 interneuron group (AVA, AVE and RIM). As a control, the neural populations in naive and trained animals displayed comparable loadings for all 3 TCs when worms were stimulated with transitions from Buffer to Buffer (Figure S8a). Together, these results demonstrate that the aversive experience with PA14 modulates population neural activity in a context-gated manner to encode learning.

To examine the specificity of PA14-induced modulation, we trained the worms with a PA14 mutant generating reduced virulence due to inactivation of a virulence factor *gacA*[36]. As we previously showed, training with PA14-*gacA(-)* did not induce a significant change in olfactory preference between PA14-*gacA(-)* and *E. coli* OP50[12] (Figure S5e,f). By analyzing the “sensory-inter group I” neural population in the discrimination task, we found that while PA14-*gacA(-)* elicited clear responses in most of the sensory neurons, training with PA14-*gacA(-)* did not significantly alter the response amplitude or the TC loadings for the population activity patterns (Figures S5c, S6c, S8b). Together, these results indicate that the neuronal and global modulations generated by the aversive training with PA14 is specific and depends on the pathogenesis of the training bacterium.

We also examined whether PA14 training-induced modulation to PA14-evoked neural responses in the discrimination task with OP50 context resulted from altered responses to OP50 by analyzing “sensory-inter group I” neurons in PA14-trained worms when stimulated with OP50 in the context of Buffer. We found that training only generated minor effects by altering the calcium signal of AWA neurons and TC3 loading of ASI (Figures S5a, S6a, S8c), consistent with comparable behavioral preference for OP50 over Buffer in both naive and PA14-trained worms (Figure S5g). Together, our results show that aversive training with PA14 modulates the activity of the nervous system in a manner that is both context-gated and task-specific. As a result, trained worms reduce their preference for PA14 in the discrimination task between PA14 and OP50, but preserve their ability in odorant detection task and continue to exhibit the preference for food over a non-food stimulus (Buffer), even if the food, PA14, is harmful.

### Learning induces context-gated modulations to different network layers

We found context-gated, learning-modulated changes in neuronal responses to PA14 in different layers of the network, including a wide range of classes of sensory neurons, interneurons, and a small number of motor neurons (Figures 1, 2, 3a). We asked whether the changes observed in the downstream neurons simply resulted from transduction of changes in neural activity at the sensory level. To test this, we fitted convolutional kernels between each pair of learning-modulated sensory neurons and learning-modulated interneurons (Figure 3b). The kernel describes what would be observed in the downstream neuron due to a simple impulse in the activity of the upstream neuron alone. While this is an abstraction and simplification, comparing the naive kernels and the learned kernels do inform us of potential changes in the network. As shown in Figure 3b, many substantial differences across the sensory-interneuron kernels suggest that it is highly unlikely that the learning-dependent changes in the interneuron activities are merely a downstream effect of the changes in the sensory neurons. Notably, by examining the kernels of ADF, ASI, AWA to downstream interneurons, we see that many of the naive kernels were negative, suggesting these sensory neurons inhibit downstream interneurons; in contrast, in the learned condition, ADF becomes less inhibitory to several downstream interneurons and ASI and AWA become slightly excitatory.

**Figure 3.**
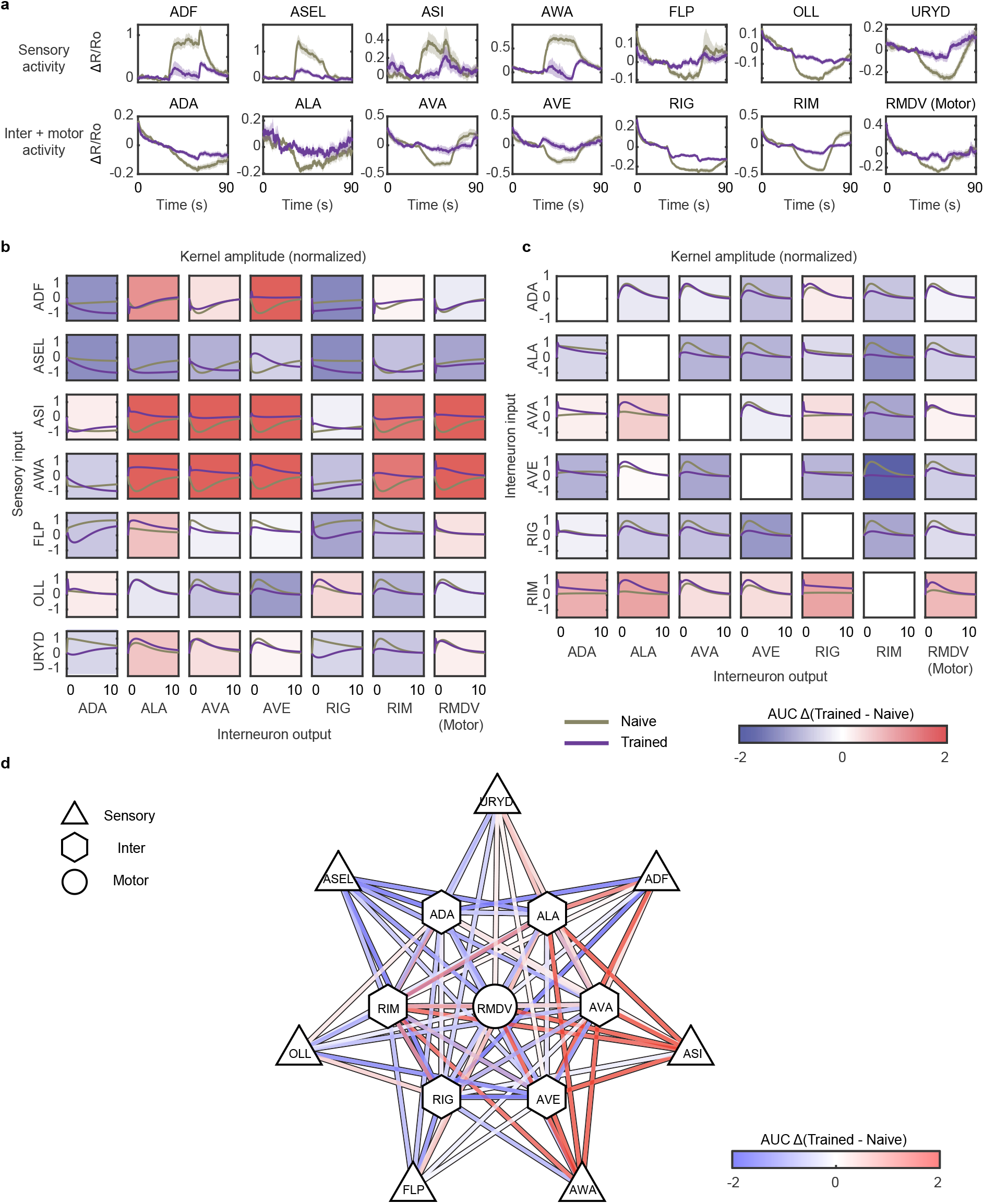
Learning modulates multiple layers of *C. elegans* neural network independently. (a) Average activity and SEM (shaded region) of neurons with significant learning-induced changes in response to OP50-PA14-OP50 stimulation pattern (naive, gray; trained, purple). (b) Double exponential kernels fitted between sensory (rows) and inter/motor neuron (columns) activities show significant divergence between naive and trained animals, indicating independently generated learning-dependent modulations to the sensory and inter/motor neuron layers. Plotted kernels correspond to averages across 2,000 bootstraps of average activation (Methods). Background color indicates normalized change in average kernel amplitude (AUC Δ) between naive and trained. (c) Convolutional kernels between interneurons also suggest changes in strength of input-output relationship between pairs of interneurons, *e*.*g*. RIM becomes less sensitive to input from other interneurons. (d) Network diagram summarizing changes in AUC between neurons from different layers.

Similarly, we also fitted kernels between the interneurons themselves to identify potential changes among the interneuron connections (Figure 3c). Consistent with the mechanisms of independent modulations, we also observed widespread changes in the kernels, further strengthening the idea that the network remodels pervasively upon learning (Figure 3d). In particular, we observe that some interneurons become less sensitive to inputs from other interneurons, e.g. AVE, RIM, and RMDV (motor). Further, ALA, AVE, and RIG drive other interneurons less strongly upon learning, demonstrated by decreased kernel amplitudes. With the exception of RIM, which drives other interneurons slightly more strongly after learning, trained kernels generally become weaker (Figure 3c), suggesting that many interneuron connections become less reliable upon learning.

To ask whether the kernel changes are context-gated, we next compared the discrimination task versus the detection task (Figure S9). On the whole, we observe more changes in the discrimination task than in the detection task. This shows that while simple sensory responses, *i*.*e*. being able to detect odorants of the pathogenic bacteria, are not altered during learning, the changes in the nervous system are much more manifested during the relevant task of bacteria discrimination. This result reinforces our notion that the changes in the nervous system upon learning are context-gated. For *C. elegans* to survive in the wild, it is adaptive for the learning to be context-specific, with changes only relevant to the task at hand and not for all interactions with the environment.

### Context-gated regulation alters network modularity

Given the widespread learning-induced changes in pairwise relationships between neurons, we next asked how these changes might alter network-level organization of the neural activities. In particular, we focus on the sensory layer, since a significant portion of sensory neurons changed their responses to PA14 after training in the discrimination task with OP50 context (Figures 1, 2). To do so, we computed modularity, an often-used metric for evaluating community structure in networks[37]. Modularity measures the degree to which neural correlations cluster into distinct modules, subsets of the network that correlate more strongly with each other than with the rest of the network. We performed such analysis on the different groups of neurons and for different stimulation patterns (Figure 4, Figures S10 and S11, and data in github repository). In particular, we found that in response to OP50-PA14-OP50 stimulation, the modularity of sensory-inter group I is relatively stable in naive animals but becomes sensitive to transitions between bacterial odorants after learning (Figure 4a). In contrast, sensory network modularity is highly responsive to Buffer-PA14-Buffer stimulation in naive animals, and only becomes slightly less responsive after learning (Figure 4b). This implies that both naive and trained animals are still sensitive to the presence and absence of PA14 bacterial odorants regardless of the training status, unlike in the bacteria discrimination task (OP50-PA14-OP50).

**Figure 4.**
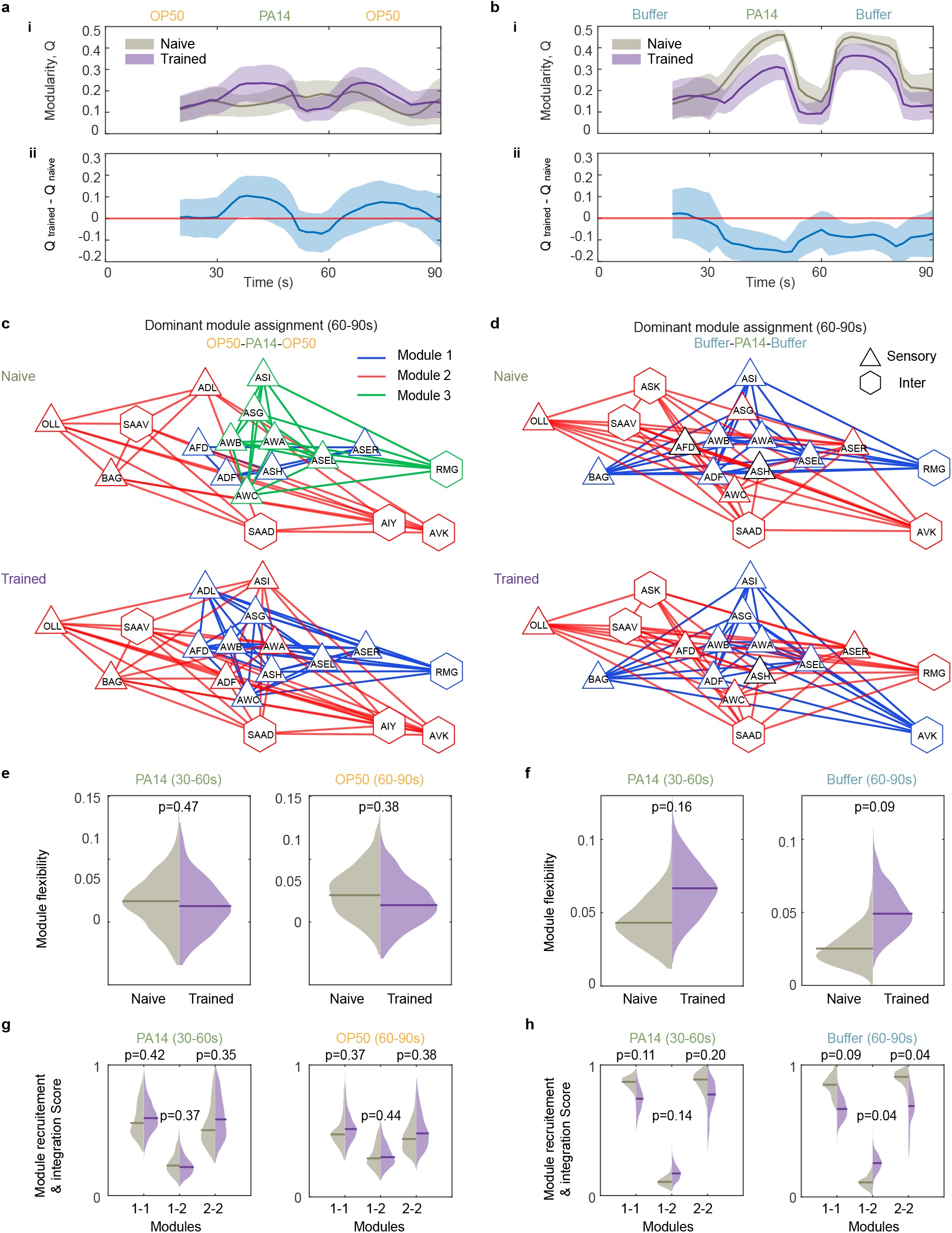
Context-gated learning-induced modulations to network-level organization of sensory neuron activities. (a, b) Modularity increases after learning in response to OP50-PA14-OP50 stimulation pattern (a, discrimination task) but decreases after learning in response to Buffer-PA14-Buffer stimulation pattern (b, detection task), suggesting increased sensitivity to differences between PA14 and OP50, but less sensitivity to PA14 alone. Both differences are consistent across bootstrapped datasets (shaded regions representing 5th – 95th percentile of bootstrapped modularity (i) and differences (ii), red line being zero difference as a guide for visualization). (c, d) While recruitment values are similar across naive and trained during discrimination between PA14 and OP50, module assignments reveal reorganization of modules after learning. Naive worms recruit three dominant modules in response to PA14 removal transitions in OP50 context (c), but trained worms recruit two dominant modules, with reorganization of module members (ADL and ASG join module 1, ADF and AWA join module 2, c). In contrast, the modules recruited by PA14 removal transitions in Buffer context remain stable (only ASH changes modules, d). (e, f) Naive and trained network flexibility are similar in response to PA14 onset and removal in OP50 context (e), but flexibility increases after learning in response to PA14 transitions in Buffer context, which is consistent with less reliable recruitment of neurons by PA14 (f). (g, h) Module recruitment and integration between modules remain similar across naive and trained during both PA14 onset (30 - 60s) and PA14 removal (60 - 90s) in OP50 context (g), but module recruitment by PA14 onset and removal transitions in Buffer context decreases after learning (and integration between modules increases). Decreased recruitment and increased integration are most notable for PA14 removal transitions, but also occur more weakly for PA14 onset transitions (h). Violin plots show distribution of values obtained from bootstrapping (n = 1,000) and median value across bootstraps (horizontal line). All p-values were obtained from bootstrapped confidence intervals.

To gain more insight into how network-level organization might be changing during learning, we inspected the underlying module assignments (Figure 4c, d, Figure S11). In response to OP50-PA14-OP50 stimulation, we found that the naive response to PA14 to OP50 transitions (60-90s) recruits three different modules of neurons (Figure 4c, top). In contrast, the trained response recruits only two modules (Figure 4c, bottom), which are the same modules recruited by OP50 to PA14 transitions (Figure S11a). This instability in naive, but increased stability of trained recruitment patterns is also shown by the stimulus-specific allegiance matrices (Figure S11c), which summarize how frequently pairs of neurons are assigned to the same module during application of these stimuli. This reorganization of network modules during learning is context-gated, as we see no such reorganization of recruitment patterns in response to Buffer-PA14-Buffer stimulus (Figure 4d, Figure S11b,d), with only ASH changing modules in this context, opposed to the OP50-PA14-OP50 condition, where several neurons change modules (ADF, ASER, AWA, ADL, ASG). In the Buffer-PA14-Buffer detection task, the decreased sensitivity of network organization to PA14 odorants seems to result from decreased recruitment, increased integration[38] (Figure 4h), and increased network flexibility[39], *i*.*e*. more frequent switching of module assignments (Figure 4f), rather than reorganization of network modules. In the OP50-PA14-OP50 context, network flexibility and module recruitment remain similar (Figure 4e, g). As a control, animals trained on PA14-*gacA(-)*, the non-virulent variant of PA14, which does not induce learning, show only small fluctuations in modularity difference from the naive animals during the discrimination task (OP50-PA14-*gacA(-)*-OP50) (Figure S10a,d,g,j,m). Two additional controls (OP50 detection task with OP50-Buffer-OP50 pattern and Buffer-Buffer-Buffer stimulation pattern) also demonstrate no appreciable differences between trained and naive animals in all modularity measures (Figure S10b,e,h,k,n, and Figure S10c,f,i,l,o). Together, these modularity results again strongly support that learning-dependent changes in network organization are also context-gated.

### Learning restructures the low-dimensional neural state manifold

We showed that across different neuron classes, the responses to PA14 in trained worms are significantly different from those in naive worms in the discrimination task. To understand how learning affects the coordination among neurons, we performed dimensionality reduction using singular value decomposition (SVD) along the neuron-axis for the combined data from the discrimination task (OP50-PA14-OP50) and detection task (Buffer-PA14-Buffer). We examined the projections of the neural activity data on the first two Neural Components (NCs) in naive and trained animals during the onset and removal of PA14 in all neuron groups (Figure 5a and Supplementary Figure S12a), and analyzed the geometry in this low-dimensional neural state[40]. We identified two types of learning-induced changes, *rotation* and *contraction*, in the neural state trajectories (Figure 5a, Supplementary Table 1 and Methods).

**Figure 5.**
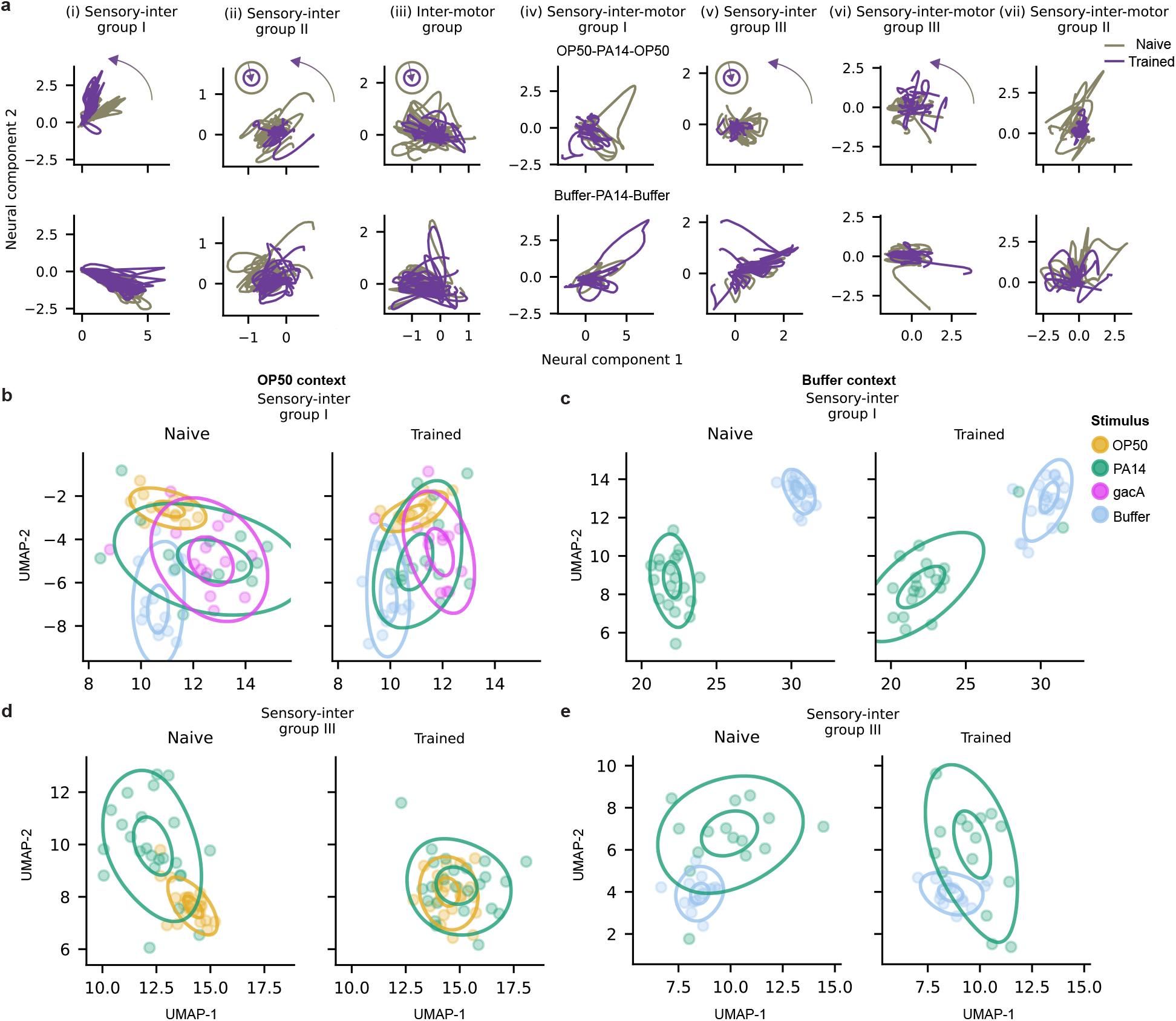
Aversive learning modulates neural state. (a) Two-dimensional neural state plotted for different neural populations (columns) in response to PA14 with OP50-PA14-OP50 stimulation pattern, top) and with Buffer-PA14-Buffer stimulation pattern (bottom). Each trace in these plots represents an individual naive (gray) or trained (purple) worm’s low-dimensional neural state. Significant rotations are marked by curved arrow and significant contractions by concentric circles with an arrow pointing to their center (Methods). (b-e) UMAP plots showing fixed-points (dots) and kernel density estimates (ellipses) for different stimuli under OP50 (b,d) and Buffer (c,e) contexts for naive (left) and trained (right) worms for sensory-inter group-I (b,c) and sensory-inter group-III (d,e). Individual data points denote fixed points for individual worms for a given stimulus with colors indicating the odor used for stimulation-PA14, gacA (PA14-*gacA(-)*), OP50 and Buffer. Inside and outside ellipses represent contours that enclose 50% and 90% of the probability mass under the 2D kernel density estimates.

First, significant rotations were detected for “sensory-inter group I, II, and III” and “sensory-inter-motor group III” neural populations (Figure 5a) revealing a strong rotating effect on sensory neurons after training and coordinated modulations of their responses. Furthermore, these rotations were specific for the PA14 and OP50 discrimination task and specific for training with the pathogenic PA14, because they were not observed for the same neuron population when PA14-trained worms were tested for the OP50-Buffer-OP50 or when worms were trained with the virulence-reduced PA14-*gacA(-)* (Figure S12b,c). Second, learning led to a contraction of the neural state trajectories in the low-dimensional space (Figure 5a). These contractions imply that the population activity deviated less along the directions of co-activity between neurons in the trained worms than in the naive worms. Significant contractions were detected in “sensory-inter group II and III” and “inter-motor group” neurons after training, uncovering a strong contracting effect on interneurons (Figure 5a). The differential training effects (rotation versus contraction) on the sensory neurons versus interneurons are surprising, suggesting that learning modulates both how interneurons process the signals from sensory neurons and/or the signals among interneurons, consistent with the results of our kernel analysis (Figure 3).

### Aversive learning directs brain-wide neural response towards predisposed configurations

Finally, we investigated how the learning-dependent global modulation of the neural responses gave rise to an activity pattern encoding learned behavior. To address this question, we examined the state of neural populations using fixed points of the neural responses evoked by different olfactory stimuli (OP50, PA14, Buffer, PA14-*gacA(-)*) and contexts (OP50 context in discrimination task and Buffer context in detection task). The presence of fixed points is suggested by the neural state trajectories in NC1 and NC2, which initially moved away from the starting point during the PA14-onset period, slowed down and stabilized as PA exposure continued, and then returned during the PA14-removal period in the discrimination task (Figure 5a). We first modeled the time series of neuronal activity for each worm and for each neural population using a Linear Dynamical Systems model[41] (Figure S13 and Methods). We then applied the validated models to identify the fixed points *x(t)*\*, where *x(t+1)*\* remained the same as *x(t)*\* [42] (Methods). Next, we compared the fixed points for neural responses to different stimuli in different contexts (Figure 5b-e, Figures S13, S14 and Methods). We found that for sensory-inter-group I, the fixed-points for the PA14 stimulation were initially coincident with the PA14-*gacA(-)* stimulation in naive worms, consistent with the similar nature of these stimuli. After training, the fixed points for PA14-evoked responses in PA14-trained worms diverged from PA14-*gacA(-)*-evoked responses in PA14-*gacA(-)*-trained animals and moved towards the Buffer fixed point (Figure 5b). This realignment of PA14 fixed point can be qualitatively observed in the 2-dimensional neural state comparing Buffer and PA14 stimulations under the OP context (Figure S12b), where the direction of the trained OP50-PA14-OP50 neural state rotates to become aligned with that of OP50-Buffer-OP50. For another 3 neuron populations (sensory-inter group II and III and inter-motor group), after training with PA14 the fixed points for PA14-evoked neural responses consistently converged onto the fixed points for OP50-evoked responses (Figure 5d, Figures S13, S14). In contrast, the fixed points for the PA14-evoked neural responses in the Buffer context for the detection task did not show consistent convergence onto the fixed points of Buffer after training (Figure 5c,e, Figures S13, S14). Thus, aversive training directs the PA14-evoked activity dynamics to a predisposed state that is expressed when the animals are exposed to no-food (Buffer) or the food that they grow on, and that the convergence of the network activity takes place in a context-gated manner to preserve the global response to PA14 in the context of no-food.

## Discussion

Several network-level strategies for encoding learning have been identified, including top-down functional remapping[5], coding by activation sequences[6], and sparse coding[2]. Among these, sparse coding explains how the nervous system integrates new information while minimizing interference with existing functions by engaging only a small subset of neurons[2]. Context-gated neural activity has also been observed in the prefrontal cortex (PFC) neurons of rhesus monkeys during working memory tasks[43–45] and in the mouse visual cortex during visual learning tested with or without the presence of a threat[46]. These findings highlight a remarkable level of flexibility for the neurons in these brain regions to respond to contextual difference with varying activity patterns.

Here, based on brain-wide functional imaging with a single-neuron resolution, we propose a new mechanism for the network-level encoding of learning: brain-wide context-gated coding that generates task-specific behavioral changes. Context-gated neural activity generates flexible neuronal responses under various external conditions, internal states and during movements[47–49]. We demonstrate, cell-by-cell and across different layers of the nervous system, that context-gated activity enables encoding of learned information that is expressed only under the context consistent with the training goal (*i*.*e*. altered olfactory preference between two bacterial smells), without interfering with the brain’s response in other contexts (*e*.*g*. preference for food smells over non-food stimuli). Interestingly, the network-level modulation by learning does not simply result from learning-induced activity changes in sensory neurons. Rather, our results suggest independent modulation of functional connectivity between sensory neurons and interneurons, as well as among interneurons, to encode learning in a context-gated manner at multiple loci of the neural network. Together, we demonstrate context-gating as a strategy that is used consistently and systematically throughout the brain to coherently encode newly learned behavior while preserving the original and essential neural functions. We further propose that brain-wide context-gated neural responses increase the capacity of information encoding by enabling flexible representations of the stimulus depending on the context. This mechanism is ecologically relevant in that it facilitates the integration of learning-dependent modulations with existing essential functions of the brain.

Catastrophic forgetting is a commonly observed phenomenon in artificial neural networks, wherein plasticity on a shared set of synapses for a subsequent task can lead to loss of prior memories[50]. This is especially a problem for small networks due to a higher probability of overlap between synapses for different memories. This may be true for the *C. elegans* nervous system, which is compact with only one to four neurons for each distinct cell type. A context-gating mechanism in combination with a reuse of neural representations could be an enticing mechanism to mitigate the challenges imposed by catastrophic forgetting for small nervous systems, which cannot afford the robustness conferred by redundant neuron types in the mammalian nervous system. Although context-gating has been previously proposed as a mechanism to attenuate catastrophic forgetting, these methods typically rely on gating which synapses to change during learning[51]. By analyzing 78% of all neuronal cell types in the head, we show that the *C. elegans* brain consistently uses context-gated neural activity as a systems strategy to encode learned information while maintaining the ability to display naive response when needed. This strategy is simple and not constrained by the degree of the redundancy of the nervous system, and thus, potentially generalizable to neural networks of other biological systems, and for artificial neural networks.

## Materials and Methods

### Strains

Adult *C. elegans* hermaphrodites were used in this study. The worms were grown and maintained according to the standard protocol at 20-22 °C in 6 cm plates containing nematode growth medium (NGM; 3 g/L NaCl, 2.5 g/L Bacto Peptone, 1.6% agar, 5 mg/L cholesterol, 25 mM *KPO*_4_ pH 6.0, 1 mM CaCl_2_, 1 mM MgSO4) seeded with the *E. coli* strain OP50 cultured with Luria Broth[52]. The *C. elegans* strains used in this study include N2 (wild-type, Bristol), ZC3292 yxEx1701[*glr-1p::GCaMP6; glr-1p::NLS-mCherry-NLS*], ZC3332 yxEx1718[*inx-4p::GCaMP6; inx-4p::NLS-mCherry-NLS*], ZC3666 yxEx1963[*inx-4p::GCaMP6; inx-4p::NLS-mCherry-NLS; mbr-1p::GCaMP6; mbr-1p::NLS-mCherry-NLS*], ZC3368 yxEx1743[*acr-5p::GCaMP6; acr-5p::NLS-mCherry-NLS*], ZC3305 yxEx1708[*ncs-1p::GCaMP6; ncs-1p::NLS-mCherry-NLS*], ZC4061 yxEx2292[*nmr-1p::GCaMP6; nmr-1p::NLS-mCherry-NLS; flp-3p::GCaMP6; flp-3p::NLS-mCherry-NLS; flp-7p::GCaMP6; flp-7p::NLS-mCherry-NLS; sro-1p::GCaMP6; sro-1p::NLS-mCherry-NLS*], ZC4164 yxEx2361[*odr-2(2b)p::GCaMP6; odr-2(2b)p::NLS-mCherry-NLS; odr-2(18)p::GCaMP6; odr-2(18)p::NLS-mCherry-NLS*]

### Generation of transgenes and transgenic animals

The DNA plasmids for multicellular calcium imaging transgenic lines were generated using Gateway LR recombination (Invitrogen #11791020) between the promoter sequence in the Gateway entry vector pCR8 (Invitrogen # K250020) and the DNA sequence for GCaMP6[20] or nuclear-localized mCherry[21] in a destination vector. To generate promoters for these plasmids, we used 5.3 kb DNA sequence upstream of the coding sequence of *glr-1* [53], 4.2 kb upstream of *acr-5* [54], 3.1 kb upstream of *ncs-1* [55], 3.5 kb upstream of *inx-4* [56], 5 kb upstream of *mbr-1* [57], 1.1 kb upstream of *nmr-1* [58], 1.5 kb upstream of *flp-3* [59], 200 bp upstream of *flp-7* [60], 1.4 kb upstream of *sro-1* [61], 2.6 kb and 2.4 kb upstream of *odr-2(2b)* and *odr-2(18)* isoforms, respectively[62]. Transgenic animals were generated by injecting transgene at 10-20 ng*/µ*l into adult animals using previously described protocols[63].

### Behavior Assays

The procedures for training worms for the droplet assay to measure olfactory learning, confocal imaging, and calcium imaging were published previously[11, 64]. *P. aeruginosa* PA14 and *E. coli* OP50 were streaked onto NGM agar plates and incubated overnight at 37 °C. On the following day, single colonies of PA14 or OP50 were inoculated into 50 ml of NGM medium and cultured overnight at 26 °C with shaking at 125 rpm for 12 hours. 1 ml of culture was evenly spread onto a 10-cm NGM agar plate to cover the entire plate[11, 65] and excess liquid was removed using a pipette. After drying the plates, they were incubated at 25-26 ° C for 36-48 hours. The training plates were placed on the bench for about 30 minutes until room temperature before use. Worms were grown on standard NGM plates seeded with OP50 until young adults. Worms at the fourth larval stage (L4) were transferred to new NGM plates with OP50 one day before the experiment. The animals were randomly picked and placed on PA14 or OP50 training plates. After 4-6 hours of training at 20 - 22 °C, worms were used for planned experiment. Training with PA14-*gacA*(-) strain and the subsequent experiments were performed using the same procedures.

### Automated droplet assay

Olfactory preference of naive and trained worms was quantified in the automated droplet setup as previously described[11, 17]. Single colonies of PA14 or OP50 were picked to inoculate 50 ml NGM medium in culture flasks and incubated overnight at 26 ° C with shaking at 125 rpm for 12 hours until the OD reached 0.3–0.35 for OP50 and 0.6-0.75 for PA14. Twelve droplets of 1.3 µl NGM buffer arranged in a 3 × 4 matrix were placed on the surface of a sapphire window that was placed in a chamber on a metal plate whose temperature was held at 22 ° C using a water bath. The testing odorants of PA14 and OP50 were delivered into the chamber using air streams that went through 20 ml of PA14 or OP50 culture contained in filtering flasks with a side tubulature (Figure S1a). Worms from training plates were first transferred into NGM Buffer (3 g/L NaCl, 25 mM KPO_4_ pH 6.0, 1 mM CaCl_2_ and 1 mM MgSO_4_) to wash briefly, then transferred into the droplets in the chamber and exposed to the PA14- or OP50-odorized air streams that alternated every 30 s for 12 cycles. The assay was recorded at 10 fps. The stimulation pattern and recording are controlled by a LabView code[11]. The locomotion of the worms was analyzed offline by custom Python and MATLAB scripts[11]. For each frame, worms were segmented from their background, binarized and fitted with an ellipse. The eccentricity timeseries of the ellipse, for each worm, was used to estimate posture and quantify turning behavior (*a* symbolizes the semi-major and *b* the semi-minor axis of the ellipse).

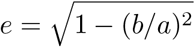

When swimming, worms occasionally bend their bodies to form a shape similar to the Greek letter Ω, which is automatically identified as a turn when the value of eccentricity falls below a set threshold. Because the frequency of Ω-shaped turns, *i*.*e*. turning rate, is negatively correlated with the preference for the stimulating odorant, a positive choice index (calculated as shown below) indicates a preference for PA14 odorants and a positive learning index indicates a reduced preference for PA14 after training[11, 17]:

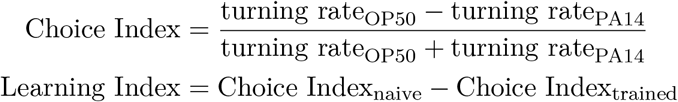

### Calcium imaging

Calcium imaging was performed in microfluidic chips by following previously established protocols[11, 66]. The microfluidic device delivered bacteria-conditioned medium or buffer to the animal’s nose and limited the Z-direction movement during recording (Figure 1a). To minimize body movement during imaging, PDMS microfluidic chips of 100% size were used for naive animals and of 90% size for trained animals. Young adult hermaphrodites were trained on *P. aeruginosa* PA14 or PA14-*gacA(-)*, or on control plate containing *E. coli* OP50 for 4–6 h before imaging, as described above. Fresh PA14 and OP50 cultures were prepared as described above for the droplet assay. Immediately before imaging, cultures were centrifuged at 3320 *× g* for 12 minutes, and the supernatants were transferred into the syringes connected with the microfluidic system. Fluorescence time-lapse imaging was performed on a Nikon Eclipse Ti-E inverted confocal microscope with a 40× oil-immersion objective and a Yokogawa CSU-X1 scanner unit and the images were recorded using an Andor iXon Ultra EMCCD camera at 5 fps. Hermaphrodite *C. elegans* were stimulated with either NGM buffer or the supernatant of a bacteria culture (*i*.*e*. bacteria-conditioned medium). Five stimulation patterns were used in this study: OP50-PA14-OP50, Buffer-PA14-Buffer, OP50-Buffer-OP50, OP50-PA14*gacA(-)*-OP50, Buffer-Buffer-Buffer, with 30-second duration for each stimulus. In each worm, we recorded time-lapse confocal images of GCaMP6 and NLS-mCherry-NLS from 1 to 4 focal planes with each focal plane containing multiple neurons (Figure 1a). The time-lapse image stacks were aligned using the StackReg plugin (rigid body transformation)[67], regions of interest (ROIs) corresponding to individual neurons were manually selected and cytoplasmic GCaMP6 and nuclear-localized mCherry signals were extracted using ImageJ. We used NLS-mCherry-NLS signals to aid in image segmentation and manual identification of regions of interest (ROIs) for quantifying cytoplasmic GCaMP6 and nuclear mCherry signals from each neuron (Figure 1 and Methods). We normalized the GCaMP6 signal from each neuron with its NLS-mCherry-NLS signal to minimize potential locomotion-related effects on signal intensity (Methods). We pooled the signals from the left and right sides of each neuron pair, except for the chemosensory neurons ASEL and ASER, which are known to be different in their activities[68]. The procedures for subsequent activity extraction and analysis are described below. Multiple worms were recorded and analyzed on separate days for each experimental condition.

### Neural activity extraction

#### Preprocessing

The fluorescence intensity of the ROIs for both GCaMP6 and mCherry channels was first background-subtracted; the ratiometric fluorescence intensity *R* was then calculated as follows:

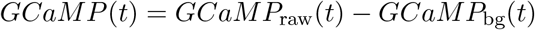

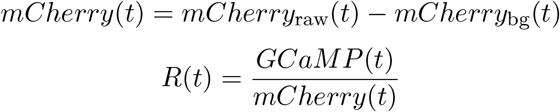

where *GCaM P*_raw_, *mCherry*_raw_, *GCaM P*_bg_, *mCherry*_bg_ are raw GCaMP6 and mCherry neuronal and background fluorescence intensity signals respectively. This was followed by baseline normalization, where the average of the last 10s of the first stimulus period of each trace was used to normalize its fluorescence intensity.

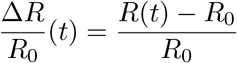

Same neuron across different focal planes was treated as technical repeats and their activity was averaged.

**Exclusion criteria** Neurons with fewer than 8 trials were excluded from further analysis.

### Confocal Imaging

Confocal Z-stack images were acquired for verifying neuron identities using colocalization of mCherry and cell-specific BFP expression (Figure S3) on a Nikon Eclipse Ti-E inverted confocal microscope with a 40× oil-immersion objective and an Andor iXon Ultra EMCCD camera. Imaging pads were prepared with 2% agar, and 3 µl of 100 mM sodium azide was applied to the agar surface. Worms were transferred into the droplet, and once immobilized, a coverglass was placed on top for imaging. Image analysis was performed using ImageJ.

### Data analysis and statistics

All code was written in Python 3 or MATLAB2021b. Dimensionality reduction, tensor operations, and imputation were performed using the tensorly libary (https://github.com/tensorly/tensorly) in Python 3.

#### Permutation test

We extracted the median neural activity for each neuron during the stimulus onset and removal periods (Figure 1a) for both naive and trained worms. Next, we shuffled the naive and trained labels 5,000 times to calculate a null distribution. We then subtracted the means of the naive from the means of the trained worms for both the data and null distributions, and calculated the chance that the absolute value of null difference was greater than the absolute value of the observed difference to get a p-value. We then applied Benjamini-Hochberg correction to the p-value to control for false discovery rate for multiple comparisons and declared significance below a threshold value of 0.05.

#### Responsive neurons

Responsive neurons were identified by comparing the neural activity flanking the odorant transition points. For each trial, there were two such points, at 30s and at 60s. For the transition at 30s, one-sample Wilcoxon signed-rank test was used to test the null hypothesis that the median of the 30-40s window was significantly different from zero. For the second transition at 60s, the median of the 30-60s window was subtracted from the 60-90s window and the null hypothesis that the signs of the transitions are symmetric about zero was tested. Benjamini-Hochberg method was applied to correct for the false discovery rate due to multiple comparisons and evaluated against a cutoff of p-value<0.05.

#### Neuron classification

Neurons were classified according to whether they encoded context, learning, or both based on the amplitude of their calcium response to PA14 (Figure 1d,e). We first tested in naive worms whether a neuron responded to the onset or removal of PA14 in either the OP50 or Buffer context (Figure S4). Neurons showing significant responses to PA14 onset or removal in either context were defined as PA14-responsive, whereas all other neurons were classified as PA14-nonresponsive. Among PA14-responsive neurons, those that responded to PA14 onset or removal in only one context, or showed responses in opposite directions in the OP50 context versus the Buffer context, were classified as context-sensitive. Neurons exhibiting similar responses to PA14 onset or removal in both contexts were classified as context-insensitive. We next compared neuronal responses between naive and trained conditions. Neurons showing significant amplitude changes during PA14-on (30-60s) or PA14-off (60-90s) periods in naive and trained worms were classified as learning-modulated, whereas those with similar amplitudes across conditions were defined as learning-independent. Furthermore, neurons showing significant learning-induced changes only in the Buffer or OP50 context were classified as context-gated, whereas those exhibiting significant learning effects in both contexts were defined as context-ungated (Figure S5h).

#### Denoising

The complete dataset was denoised using a standard wavelet denoising procedure using Daubechies-4 (db4 wavelet) up to level 4 with symmetric boundary handling before dimensionality reduction for the temporal and neural components and fixed-point finding. We estimated noise level using median absolute deviation of the coefficients at level 4 and soft threshold 1 using the *pywavelets* (https://github.com/PyWavelets/pywt) package in Python.

#### Imputation

In order to fill gaps (*i*.*e*. missing neurons) in worms for downstream analysis, we used PARAFAC (parallel factor analysis) tensor decomposition based imputation that utilizes the structure of the data to fill up missing values using *tensorly*. PARAFAC imputes the missing trials by making use of correlations throughout the whole tensor. The following criterion to do the imputations was used: Minimum number of trials of a neuron across worms to be considered: 8. Minimum fraction of neurons that could be recorded out of the total number of neurons expressing GCaMP6 for the specific transgenic strain in a worm to be considered: 2/3. Imputed neurons were masked out for dimensionality reduction.

#### Temporal components

We stacked the activity traces of all neurons from both naive and trained animals probed with the bacteria discrimination task (OP50–PA14–OP50) or the bacteria detection task (Buffer–PA14–Buffer) to get a trial time x (worms x neurons) matrix. We then extracted the data from the last 60 seconds for each trial from this matrix, i.e., without using the neural activity during the time of the anchor odorant stimulation for the decomposition. We then scaled the responses across all neurons with the feature-wise (across neurons and worms for each time point) standard deviation to bring the responses to a common scale. Temporal components were then extracted from the concatenated data with Singular Value Decomposition (SVD) using the *tensorly* library. We selected the top 3 temporal components for further analysis.

### Behavioral Components

The droplet assay involves 12 cycles of 30s alternating odorant stimulations to the worms while they are video recorded at 10Hz. This gives 12 trials of alternating anchor and contrast odorant stimulation exposure to the worms. Using a moving window of 90s and stride length of 60s, we ran through and stacked together every trial for naive and trained worms for a given pair of odorants. We then normalized the eccentricity values per trial in the last 10s of the anchor period by mean subtraction and scaling. Then, we took the last 60 seconds of this matrix, scaled by the standard deviation and performed PCA using *sklearn* library (https://github.com/scikit-learn/scikit-learn) to get the behavioral components.

To align behavioral components (BCs) with temporal components (TCs), we subsampled the behavioral components by picking every second point to match the TCs.

### Fitting convolutional kernels

We fitted double exponential convolutional kernels 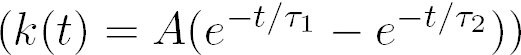 between average activity of input (sensory or inter) and output (inter or motor) neurons. Kernels were only fitted between neurons showing significant learning-dependent changes in response to OP50-PA14-OP50 stimulus (sensory: ADF, ASEL, ASI, AWA, FLP, OLL, URYD; inter: ADA, ALA, AVA, AVE, RIG, RIM; motor: RMDV). Kernel parameters (*A, τ*_1_, *τ*_2_) were fitted by calculating a discrete kernel of 150 time points (30s), convolving the discrete kernel with input neuron activity (450 time points), and least squares minimization of the resulting error between predicted and observed activity of the output neuron. To estimate the effect of variability in input and output neuron activity on kernel shape, average input and output neuron activity was bootstrapped (n = 2,000 bootstraps) for each neuron pair and kernel fitting was repeated for each bootstrap. For visualization in Figure 3, fitted kernels were averaged across bootstraps and these average kernels for each neuron pair were normalized by the maximum absolute amplitude across naive and trained conditions. This normalization preserved the relative magnitude of naive and trained kernels. To compare naive and trained kernels, we also calculated the average kernel amplitude across time (*AUC*) for each bootstrap, subtracted average naive *AUC* from trained *AUC*, and normalized by the mean absolute value of naive and trained *AUC*. This analysis was performed using MATLAB 2021b (code will be made available upon acceptance).

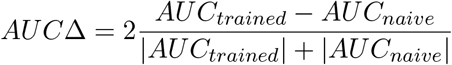

### Network modularity analysis

#### Population averaging correlations and bootstrapping

Data for each combination of imaging strain, training, and stimulus condition can be thought of as a partially observed 3D tensor (neurons x recordings x time points), where each recording contains a different combination of neurons observed across all time points. To obtain population-averaged time-varying functional connectivity, we expanded the neurons dimension into sliding window correlations (20s long [100 frames], 2s offset [10 frames]) for each neuron pair (neuron pairs x recordings x time windows), then averaged across observations (recordings), resulting in a 2D tensor (neuron pairs x sliding windows) that can be reshaped into a 3D tensor of time-varying correlation matrices (neurons x neurons x time windows). To estimate the effect of recording-to-recording variability in observed neurons and correlations on our results, we bootstrapped population averaged correlations by resampling the recordings dimension of our data tensor before averaging correlations (n = 1,000 bootstraps).

To ensure comparability of naive and trained correlation matrices, we masked out any neuron pairs with fewer than three observations in either condition for each strain and stimulus combination. This ensured both correlation matrices (within each imaging strain and stimulus combination) contained the same combination of observed and unobserved correlations. We repeated this masking procedure for each bootstrap because different recordings contained different combinations of neurons, leading to different numbers of observation per pair, even if the number of recordings per bootstrap was conserved.

#### Obtaining representative module assignments and calculating network modularity

For each bootstrap, sliding window correlation matrices, *A*(*t*), were Fisher transformed, *Z*(*t*) = *arctanh*(*A*(*t*)), and diagonal ones were zeroed out before calculating multi-slice modularity [69] and performing Louvain optimization[70, 71]. In the equation below, *i* and *j* are neuron indices, *s* and *r* are sliding window indices, *δ* is the Kronecker delta function, *g*_*is*_ is the module assignment of neuron *i* in window *s*, and *µ* is a normalization constant that sums all possible contributions to modularity correlations and coupling across windows. The 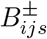 terms are the contributions of individual neuron pairs *ij* to multi-slice modularity for window *s* and incorporate a modification of modularity that accounts for correlation signs[72]. This modification split positive and negative Fisher transformed correlations 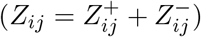 to account for signs of correlations between neurons and compares observed correlation strength 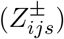, to average strength of positive/negative correlations for each neuron 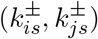 and overall strength of positive/negative correlations 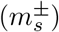 for a given window *s*.

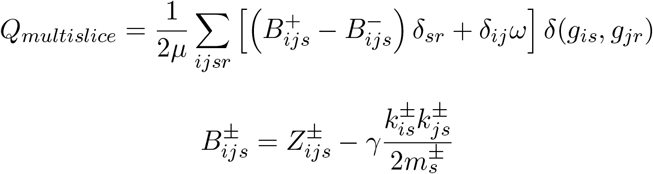

For the mutli-slice coupling parameter *ω*, we selected a value of 0.4, but observed similar module assignments and changes in modularity over time with different values (0.2 - 0.6). For the resolution parameter, *γ*, we used a value of 1.

To obtain a representative module assignment based on multi-slice modularity, we repeated Louvain optimization 1,000 times and found a representative consensus assignment across optimization runs[73]. Module assignments for each window were re-indexed to minimize the number of neurons that change module assignment across adjacent window. In addition to finding a representative consensus assignment across optimization iterations for each bootstrap, we also found a bootstrap averaged module assignment using the same representative consensus approach, which is used for visualization in Figure 4c,d and Figures S10 and S11.

For each bootstrap, we calculated modularity for each sliding window using the formulation of modularity below, which accounts for correlation signs. In the equation below, *i, j* correspond to neuron indices and *t* to sliding window indices.

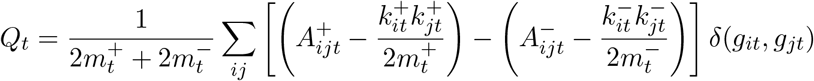

This analysis made use of the GenLouvain toolbox for MATLAB [71] and was performed in version 2021b. All code will be made available upon acceptance.

#### Calculation of recruitment and integration

Recruitment and integration were calculated according to the definitions provided by Bassett *et. al*. 2015[38]. First, the time-varying module assignments were used to calculate an allegiance matrix, *P*_*ij*_, which is the probability of two neurons being assigned to the same module across time points. Allegiance matrices were calculated for specific time periods to obtain stimulus specific recruitment and integration.

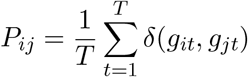

The recruitment and integration scores (*R*_*a*_ and *I*_*ab*_) for specific stimulus periods are block-wise averages of the corresponding allegiance matrix *P*_*ij*_ based on time-averaged module assignments *C*_*a*_, *C*_*b*_, *C*_*c*_, …*C*_*K*_. Recruitment corresponds to averaging within a time-averaged module (*a* = *b*)and integration corresponds to averaging across modules (*a≠ b*).

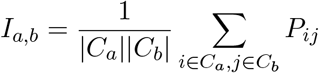

Allegiance matrices and recruitment and integration were recalculated for individual bootstraps using the same time-averaged module assignments, which were derived from consensus clustering the bootstrap-averaged module assignment.

#### Calculation of network flexibility

Network flexibility *F* was calculated according to the definitions from Bassett et. al. 2011[39]. Network flexibility is the total number of module assignments switches across adjacent windows, divided by the total number of changes possible (number of windows *T* − 1 times number of neurons *N*).

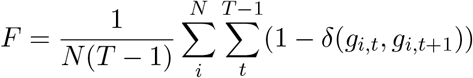

This flexibility calculation was repeated for each bootstrap of population-averaged sliding window correlations.

### Network modeling and analysis

Data, network structure and dynamics were analyzed and models were simulated using the CeDNe framework ([74]), which integrates connectomic, anatomical and functional data into a graph-based unified modeling environment. CeDNe is available online at https://github.com/sahilm89/CeDNe.git.

#### Neural components

Last 60 seconds of the trials from neurons common to naive and trained animals were stacked for each neuron group across OP50-PA14-OP50 and Buffer-PA14-Buffer stimulations along the direction of neurons without using the time of the anchor-odorant stimulation to get a neuron x (worms x time) matrix. The responses were then scaled by standard deviation and Neural Components were extracted using Singular Value Decomposition (SVD) across the neuron axis. 7 sets of neural components for the 7 neuron groups were thus produced, out of which the first two neural components from each group were used for further analysis.

#### Neural state analysis

The geometry of neural state trajectories was analyzed to quantify the changes in the low-dimensional manifold over learning. Rotations and contractions in the 2-dimensional neural state manifold were calculated by taking epochs of 30 frames (6 seconds) and sampling over the length of the trajectory in strides of 12 seconds for each worm within each neuron group, learning and stimulation conditions. Then, the average of the worms during each epoch was taken to get a epoch-averaged sequence of points in the low-dimensional neural space.

##### Contractions

Contraction was calculated by measuring the relative area of these epoch-averaged trajectories in low-dimensional space. First, area was calculated by measuring the square-root of the determinant of the covariance matrix of the neural state for each epoch in 2 dimensions and calculating the average across *n* epochs.

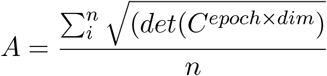

Then the trained area was divided by the naive area to get a contraction *C*.

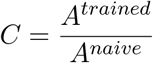

Areas of naive and trained worms were statistically compared using Mann Whitney U-test with multiple comparisons correction (Benjamini-Hochberg) with a p value-cutoff of 0.05.

##### Procrustes alignment

Rotational changes to the neural state manifold were systematically characterized over learning by performing Procrustes alignment of the epoch-averaged naive and trained trajectories to extract the set of linear transformations that maximally overlap the naive and the trained neural manifold by minimizing *d*_*proc*_.

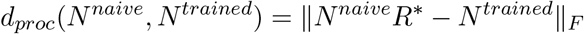

where *R* is the optimal rotation matrix, *N* ^*naive*^ and *N* ^*trained*^ are naive and trained neural state vectors and *F* is the Frobenius norm. Statistical significance of this rotation was tested by comparing against a null distribution constructed by shuffling the naive and trained labels and using a cutoff of p<0.05.

#### Linear Dynamical System

Inputs and dynamics of the data were analyzed by fitting the data to a Linear Dynamical Systems (LDS) model. LDS model was first fit to the temporal components and then later expanded to the whole neural activity data of a worm within each neuron group, stimulation and learning conditions.

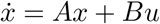

##### Temporal component fitting

For temporal components, this model was fit with ridge penalty for *A, B* at 1e-3 and L2 penalty at 1e-3. Here *x* is a vector of the 3 temporal components, *A* captures the interactions between the components, and *B* is the scaling factor for the inputs *u*, which are step-function stimulus input vectors that capture the time-courses for the 2 different input patterns (30s ON, 30s OFF and 30s OFF, 30s ON) for the last 60 seconds of anchor and contrast odorant exposures. Then the *τ* for each component was calculated from the eigenvalues of the A matrix. For each *TC*_*i*_

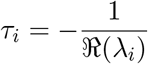

where *λ*_*i*_ is the *i*^*th*^ eigenvalue of the A matrix.

##### Fixed-point finding

For the fixed-point finding, the model above was fitted subject to the denoised, imputed, complete blocks of neural activity data for each worm with ridge parameters for *A* and *B* at 1e-1 each. Here *x* is neural activity, *A* captures the effective connectivity between neurons, and *B* is the scaling factor for the inputs *u*, which are step-function stimulus input vectors that capture the time-courses for the different odor stimulation inputs.

Fixed points were then calculated *x*^∗^ using the fitted *A* and *B* matrices by finding the steady-state:

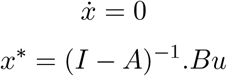

Fixed points for the 2nd and 3rd odor presentation (last 60s) were used for further analysis.

##### Contexts in fixed-point analysis

Datasets were divided based on the context odorants into 2 sets: OP50-context and Buffer-context based on the first (anchor) odorant in the stimulation paradigm to keep all odorant comparisons within the same context space. OP50-context includes OP50-PA14-OP50, OP50-Buffer-OP50 and OP50-PA4-*gacA(-)*-OP50 stimulation patterns. Buffer-context includes Buffer-PA14-Buffer and Buffer-Buffer-Buffer stimulation patterns.

## Data Availability

All data collected and analyzed in this study are included in Results, Supplementary materials, and the associated online resources. Imaging data, codes developed and applied in this study will become accessible upon acceptance.

## Acknowledgments

We thank *Caenorhabditis Genetics Center* for strains (funded by NIH Office of Research Infrastructure Programs P40 OD010440), Drs. C. Bargmann, O. Hobert, N. Ringstand, E. Yemini for strains and transgenes. H.L and Y.Z are funded by  NIH (MH130064 and NS115484).

## Author Information

### Contributions

Project conception, experimental design and result interpretation: J.L., S.Moon, S.Moza, H.J.L., P.E.E., H.L., Y.Z.

Transgene and transgenic strain construction: J.L., J.C., M.C.

Calcium imaging data collection: J.L., P.E.E., M.G.

Naive and PA14-trained strains: J.L., M.G.

Naive and PA14-*gacA(-)* trained strains: P.E.E.

Calcium imaging segmentation, cell identification, signal extractions, computations and analysis: J.L., S.Moon, S.Moza, H.J.L., P.E.E.

Online resources: S.Moon (https://github.gatech.edu/Lu-Fluidics-Lab/context-gated-learning-analysis)

S.Moza (https://github.com/Yun-Zhang-Lab/whole-brain-learning)

Code development, computational models, and application: S.Moon, S.Moza, P.E.E.

Manuscript writing: J.L., S.Moon, S.Moza, H.J.L., P.E.E., H.L., Y.Z.

Figure generation and preparation:

J.L. (Figure 1, Supplementary Figure S1, S2, S3, S5)

S.Moon (Figures 3, 4, Supplementary Figures S9, S10, S11)

S.Moza (Figures 1, 2, 5, Supplementary Figures S4, S5, S6, S8, S12, S13, S14)

H.J.L. (Figures 3, 4, Supplementary Figures S9, S10, S11)

P.E.E. (Supplementary Figure S7)

## Supplementary materials

**Figure S1.**
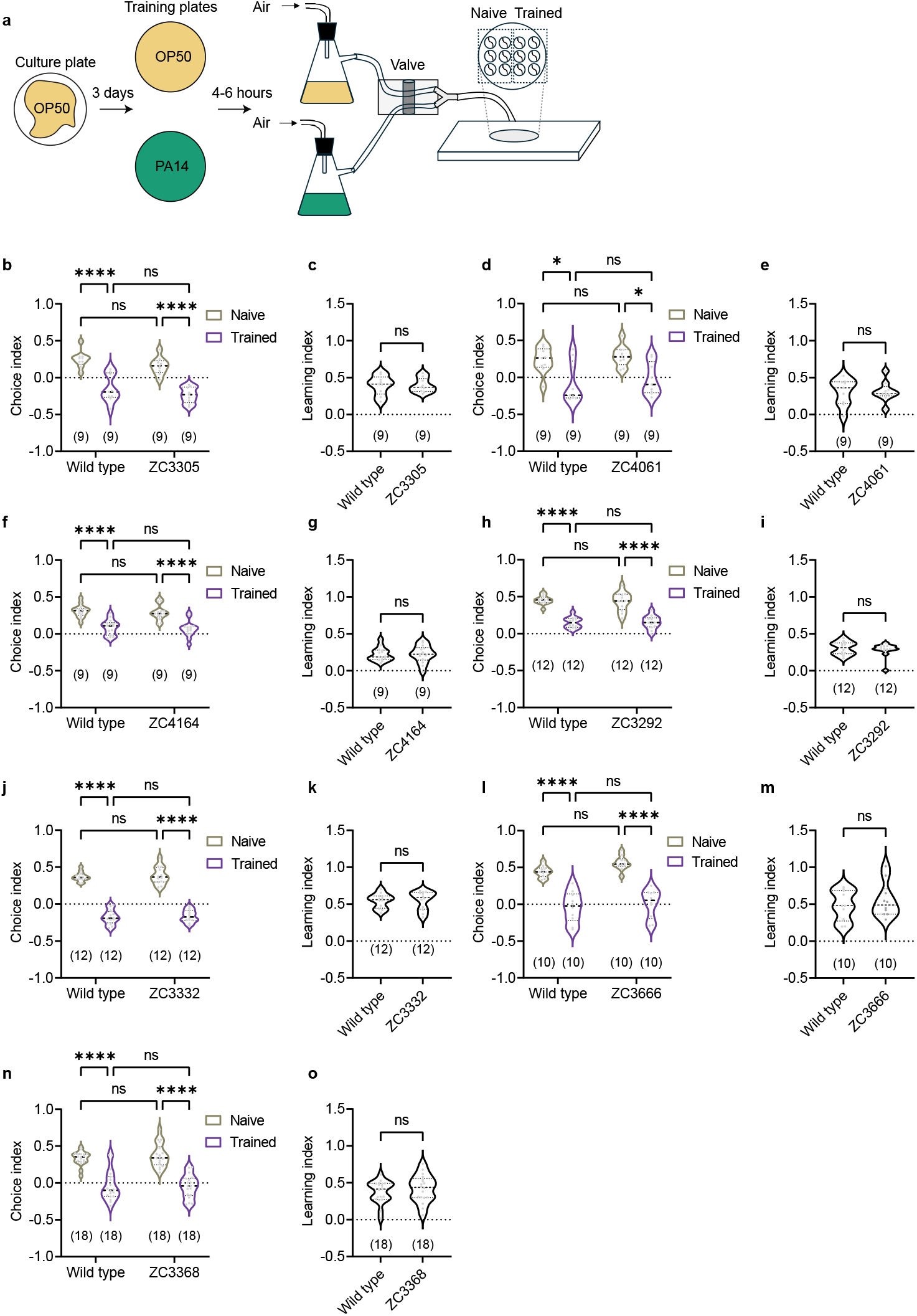
Multicellular calcium imaging strains exhibit normal learning ability. (a) Schematic of the droplet assay used to measure odorant preference. Worms were trained for 4–6 hours on plates seeded with either *E. coli* OP50 or *P. aeruginosa* PA14 before testing. (b–o) Violin plots showing choice indices between PA14 odorants and OP50 odorants or learning index for each multineuronal calcium imaging strain (see Supplementary Figures 2 and 3 for expression patterns). Violin plots illustrate the frequency distribution of data points, with the width representing the density of observations. Horizontal lines denote the median (dashed line) and quartiles (dotted lines). The number of assays is shown in parentheses, and the results of individual assays are shown as gray dots. Statistical analyses were performed using two-way ANOVA with Tukey’s multiple comparisons test for choice index and Student’s *t*-test for the learning index. Asterisks indicate significant differences: **** *p <* 0.0001, * *p <* 0.05, ns: not significant.

**Figure S2.**
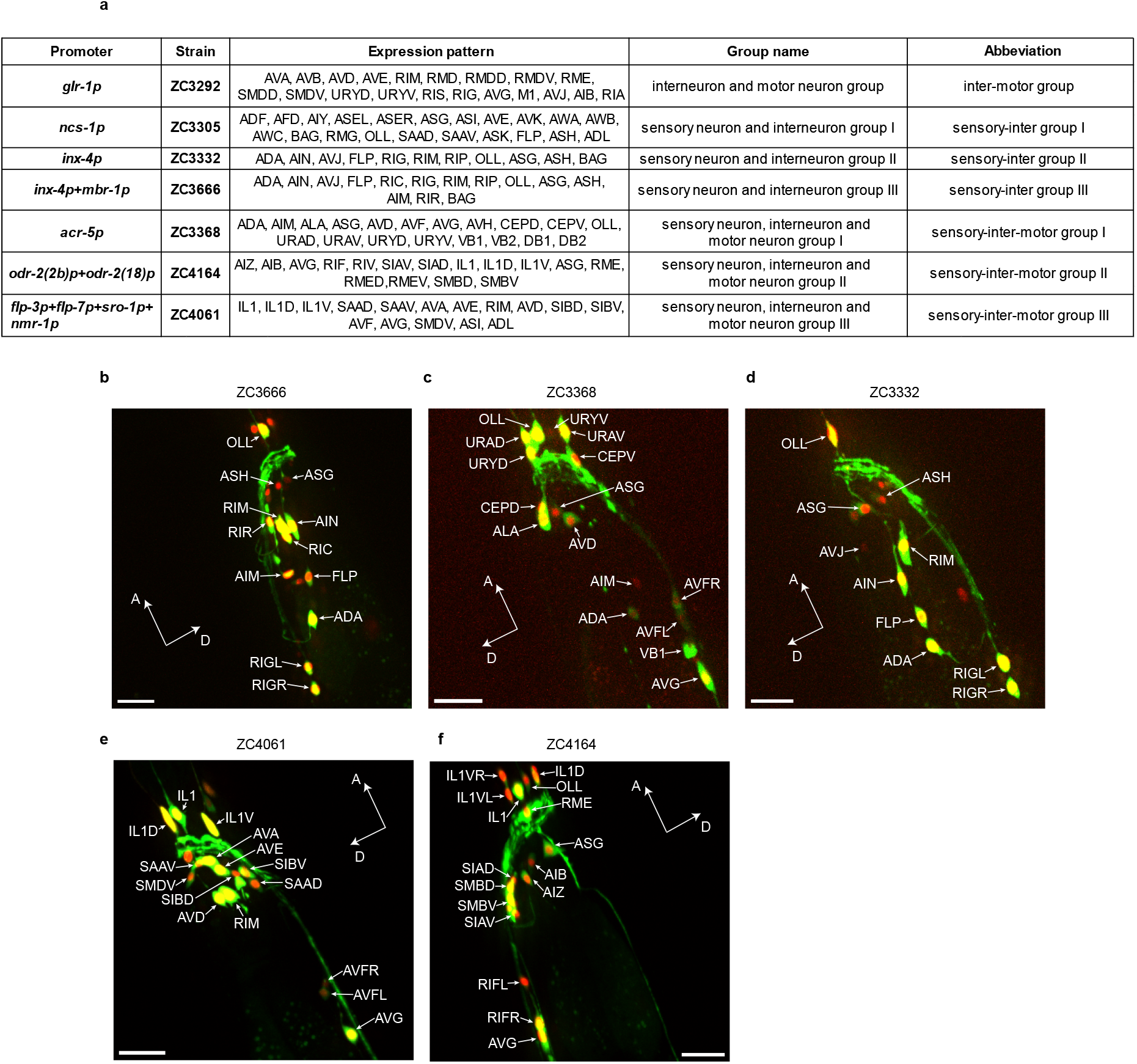
Multicellular imaging strains sparsely express cytoplasmic GCaMP6 and nuclear-localized mCherry in neuronal groups. (a) Information table for the seven multicellular calcium imaging strains used in this study. (b) Sample images for five multicellular calcium imaging strains in addition to another two shown in Figure 1c. Only one lateral side of the worm is shown. Arrowheads indicate the soma of each identified neuron class. A, anterior;D, dorsal. Scale bar, 20 *µ*m.

**Figure S3.**
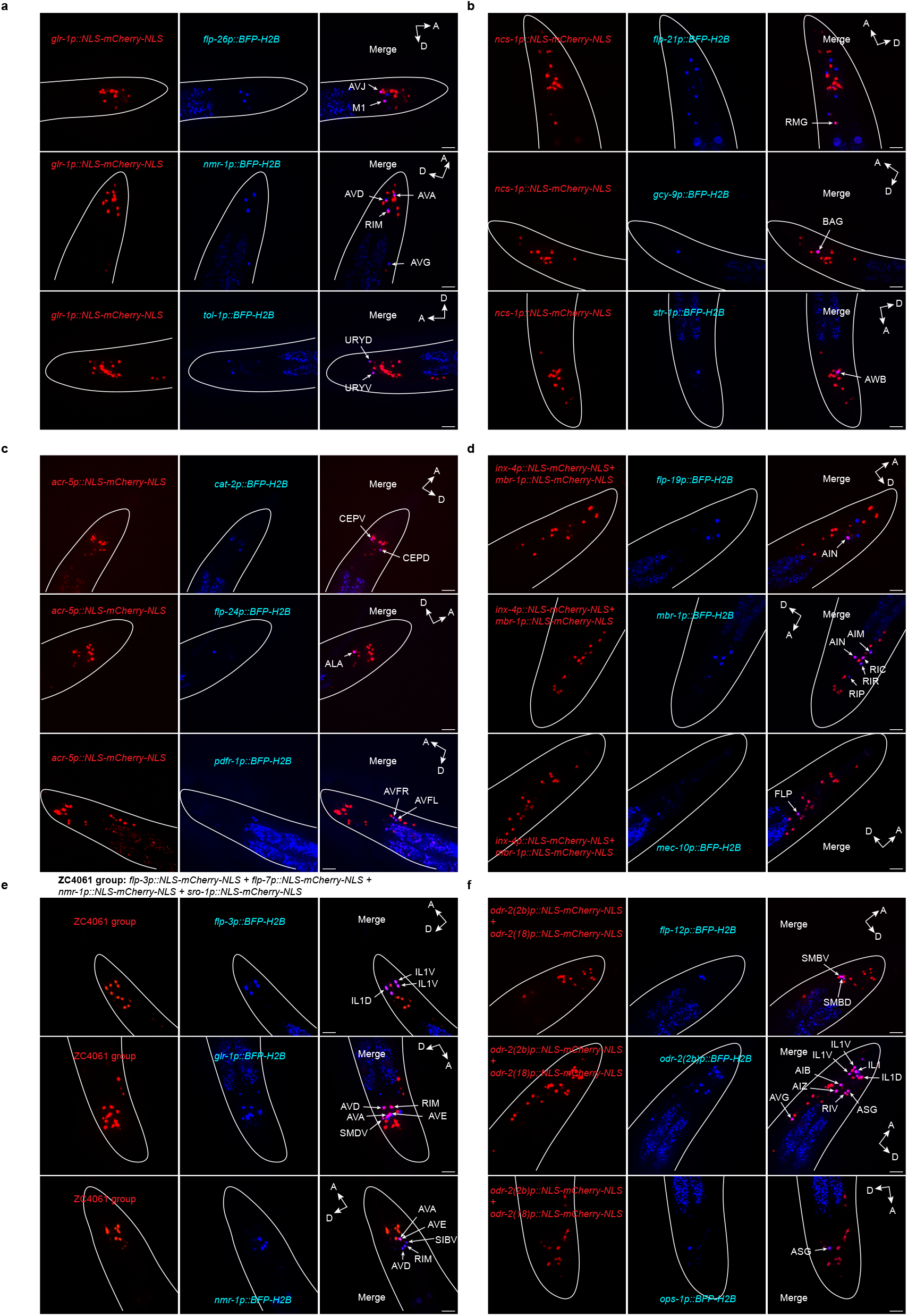
Cell-identification in multicellular imaging strains with cell-specific nuclear-localized BFP reporters. (a–f) Sample images showing co-localization of *flp-26p::BFP-H2B, nmr-1p::BFP-H2B, tol-1p::BFP-H2B, cat-2p::BFP-H2B, flp-24p::BFP-H2B, pdfr-1p::BFP-H2B, flp-3p::BFP-H2B, glr-1p::BFP-H2B, flp-21p::BFP-H2B, gcy-9p::BFP-H2B, str-1p::BFP-H2B, flp-19p::BFP-H2B, mbr-1p::BFP-H2B, mec-10p::BFP-H2B, flp-12p::BFP-H2B*, and *odr-2(2b)p::BFP-H2B* with NLS-mCherry-NLS expression in naive adult hermaphrodites of multicellular imaging strains. Arrows indicate neurons co-expressing BFP and NLS-mCherry-NLS, which were used for cell type identification in calcium imaging. Only one lateral side of the worm is shown. Worm body outlines are approximately shown with white lines. Scale bar, 20 *µ*m.

**Figure S4.**
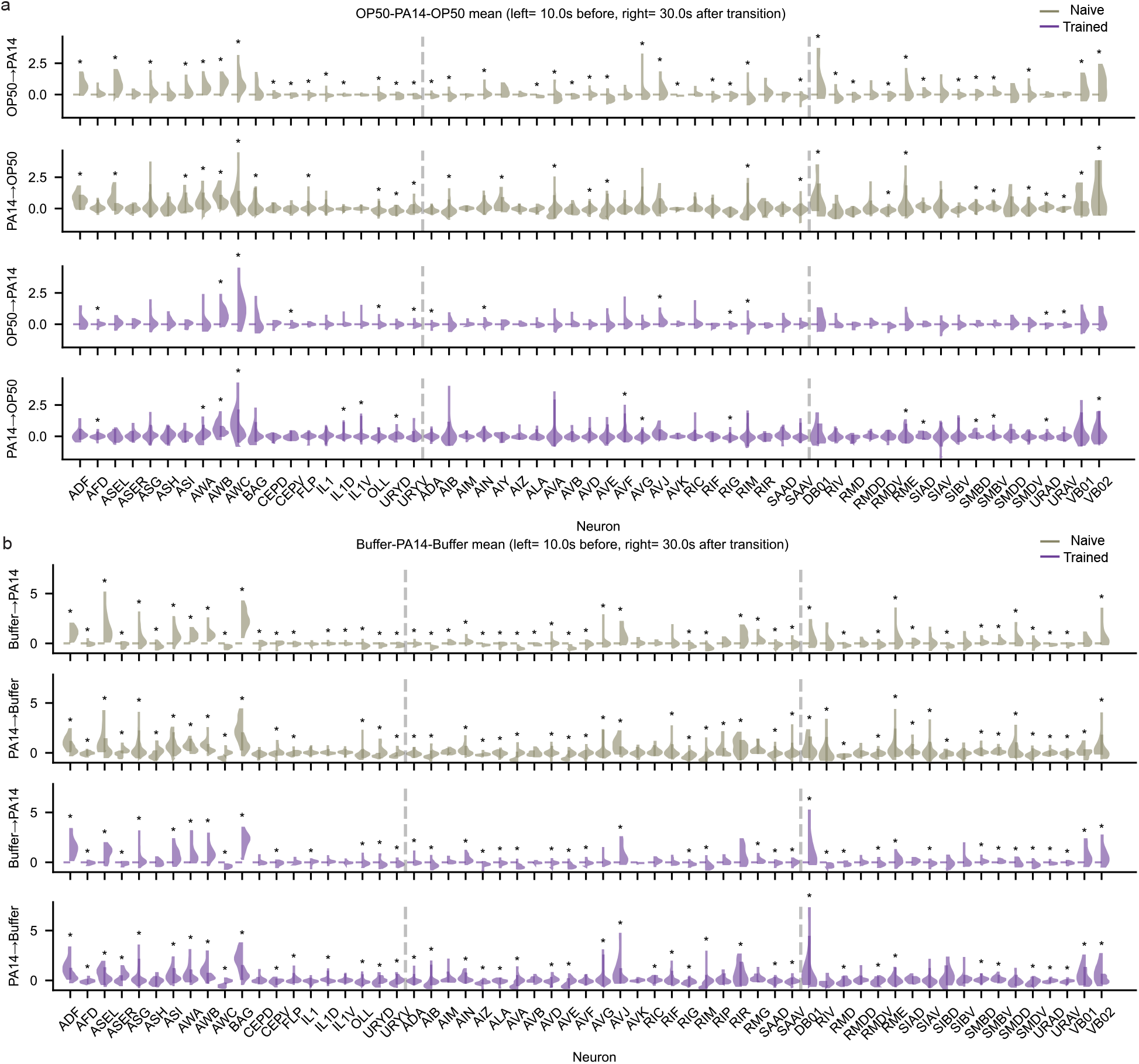
Wide-spread responses to PA14-conditioned medium in the *C. elegans* nervous system in OP50 context and Buffer context. (a, b) Violin-plots showing kernel densities of the distributions of medians for neurons in response to to PA14 transitions in naive and trained worms under OP50 context for OP50 → PA14 and PA14 → OP50 transitions (a), and Buffer context for Buffer → PA14, PA14 → Buffer transitions (b). Asterisks indicate “responsive neurons” that are statistically significantly different between the windows flanking each transition (Methods).

**Figure S5.**
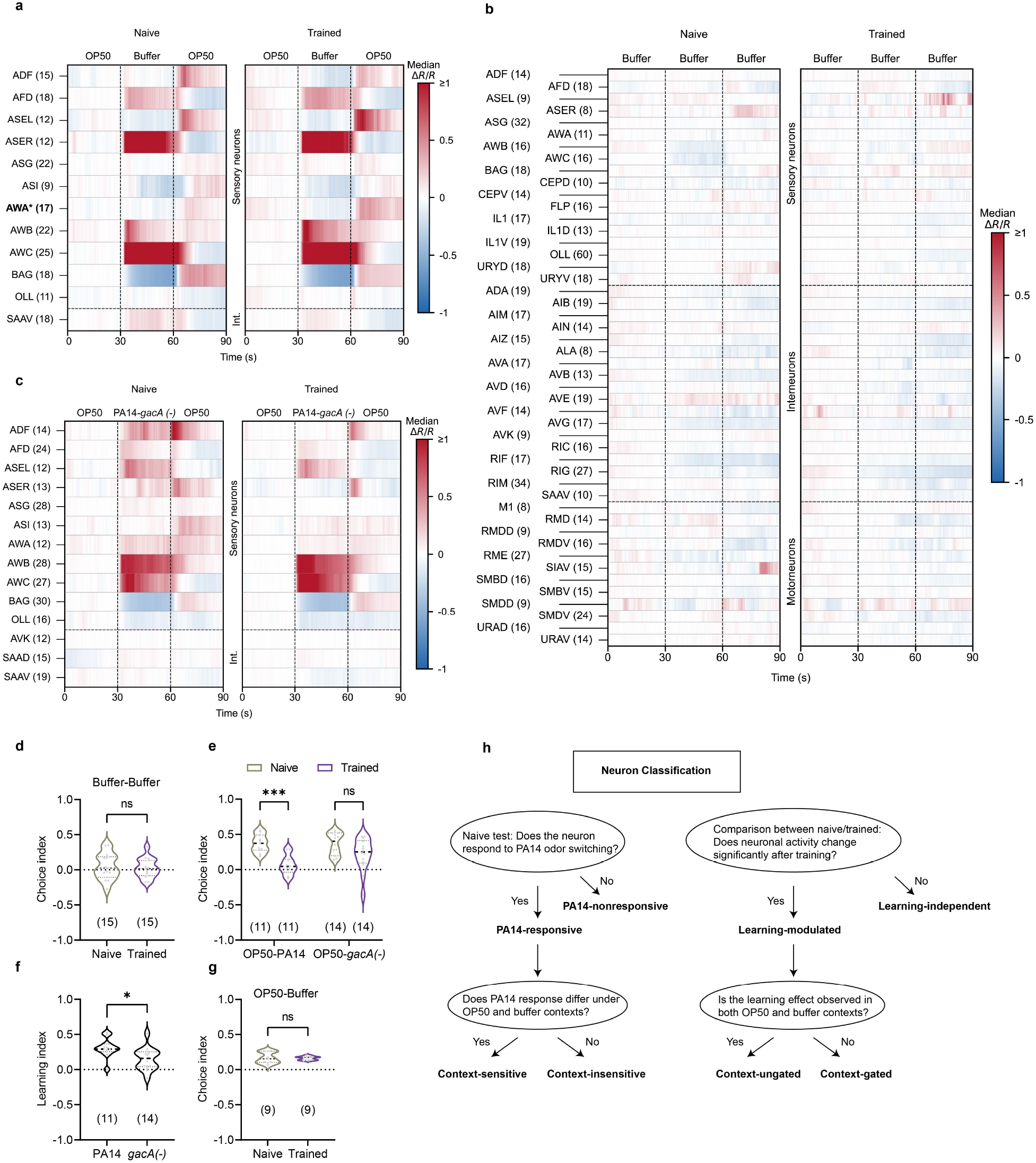
Training with PA14 induces brain-wide neuronal changes that are specific to the training bacterium and depend on its pathogenesis. (a–c) Heatmaps for neural activities measured by GCaMP6 signals in identified head neuron classes in response to OP50-Buffer-OP50 stimulation pattern in naive and PA14-trained worms (a), Buffer-Buffer-Buffer control stimulation pattern in naive and PA14-trained worms (b), and OP50-PA14-*gacA(-)*-OP50 stimulation pattern in naive and PA14-*gacA(-)*-trained worms (c). Neuron classes recorded in ≥ 8 worms are shown. * and bold font indicate significantly different neural activities between naive and trained worms in response to either the removal or the onset of OP50 (a), the transitions between Buffer (b), or the onset/removal of PA14-*gacA(-)* (c), p *<* 0.05, based on label-shuffling permutation test with 5000 iterations followed by Benjamini–Hochberg false-discovery-rate correction using median of the traces. The number in parentheses indicates the smaller number of worms recorded for that neuron class under naive and trained conditions. (d,g) Violin plots of choice indices for Buffer transitions (d), and for OP50 and Buffer preference (g) in naive and PA14-trained worms tested in droplet assays. (e,f) Violin plots of choice indices (e) and learning indices (f) for OP50 and PA14 preference in naive and PA14-trained worms (left) or for OP50 and PA14-*gacA(-)* preference in naive and PA14-*gacA(-)*-trained worms (right). Violin plots illustrate frequency distribution of data points, with width representing density of observations. Horizontal lines denote median (dashed) and quartiles (dotted). The number of assays is shown in parentheses. Statistical analyses were performed using Student’s *t*-test (d, f, g) and two-way ANOVA with Tukey’s multiple-comparisons test (e). Asterisks indicate significant differences: *** *p <* 0.001, * *p <* 0.05, ns: not significant. (h) Schematic diagram showing the workflow for neuron classification, as in Figure 1g, based on neural responses shown in Figure 1d–e.

**Figure S6.**
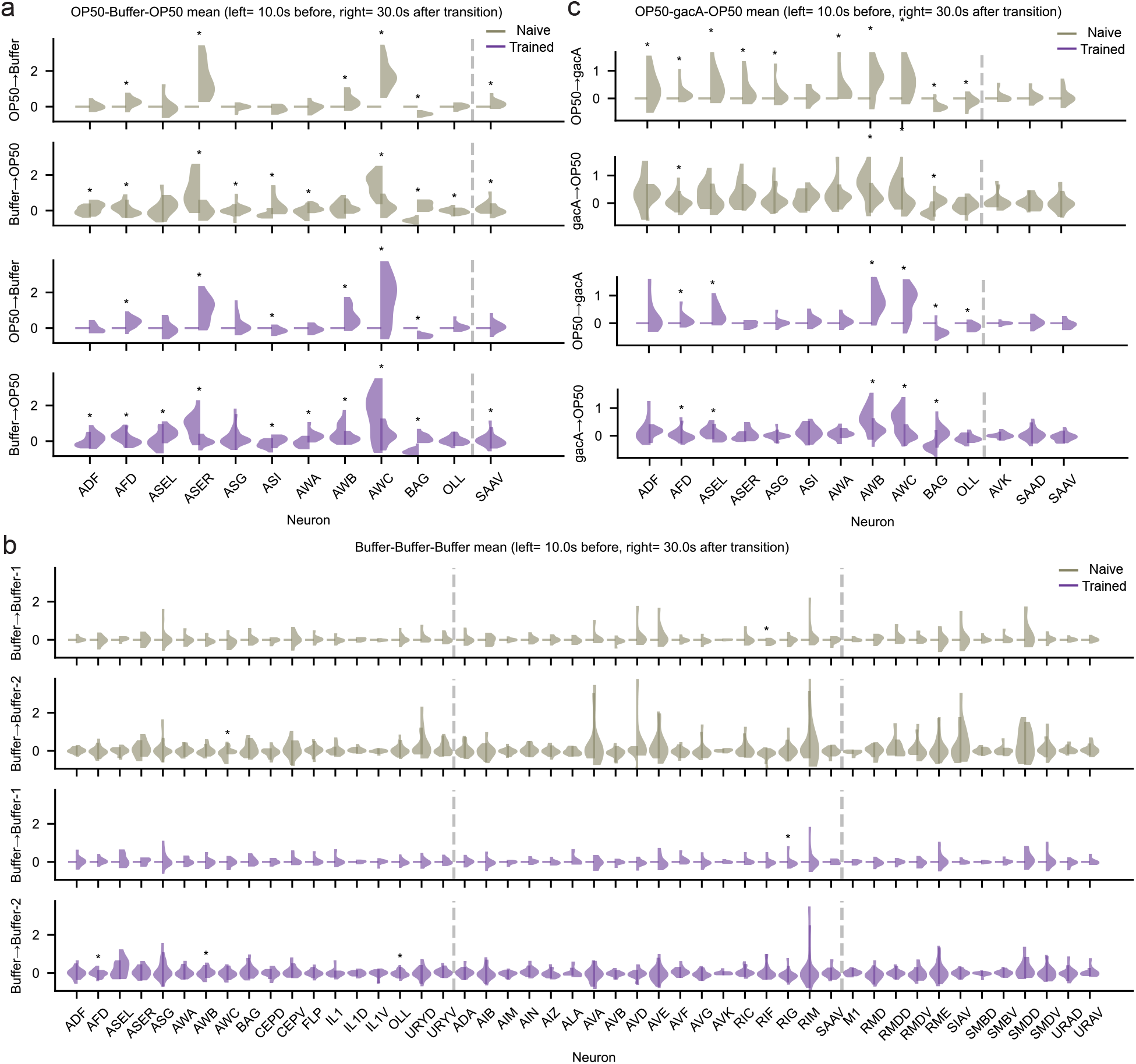
Responses to OP50-Buffer-OP50 stimulation pattern and to PA14-*gacA(-)* (referred to as gacA in this figure) in OP50 context in sensory-inter group I neurons, as well as control responses in the *C. elegans* nervous system. (a-c) Violin plots showing distribution of medians for neurons responsive to several stimulus shifts: OP50 Buffer and Buffer → OP50 transitions in OP50-Buffer-OP50 stimulation pattern (a), Buffer → Buffer transitions in Buffer-Buffer-Buffer stimulation pattern (b), OP50 → PA14-*gacA(-)* and PA14-*gacA(-)*→ OP50 transitions in OP50-PA14-*gacA(-)*-OP50 stimulation pattern (c). Asterisks indicate responsive neurons that were statistically different between the windows flanking each transition (Methods).

**Figure S7.**
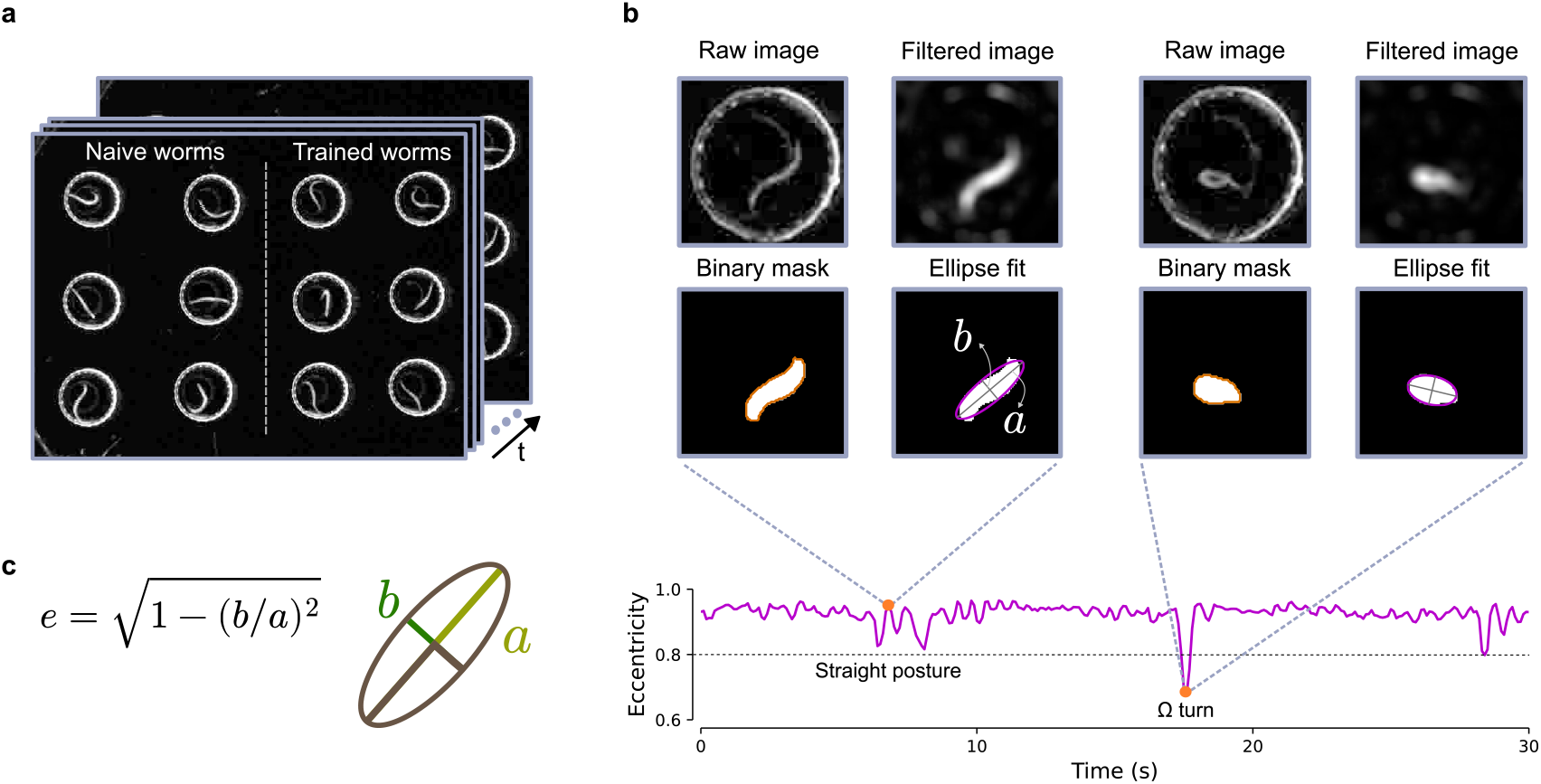
Locomotory gait quantification in swimming worms through shape tracking. (a) Video recording of naive (left) and trained (right) worms placed in droplets and exposed to sequences of odorants. (b) Frame sequences of individual worms are computationally isolated, background subtracted, filtered by blurring and edge attenuation and binarized to create clean worm masks. An ellipse is fitted to the binary mask in each frame and its eccentricity is measured over time as a proxy for worm posture (b: semi-minor axis, a: semi-major axis). A manually verified threshold for eccentricity is set, and peaks below it are considered turns. (c) Formula for eccentricity calculation where b is the semi-minor axis and a is the semi-major axis of the fitted ellipse (see also b).

**Figure S8.**
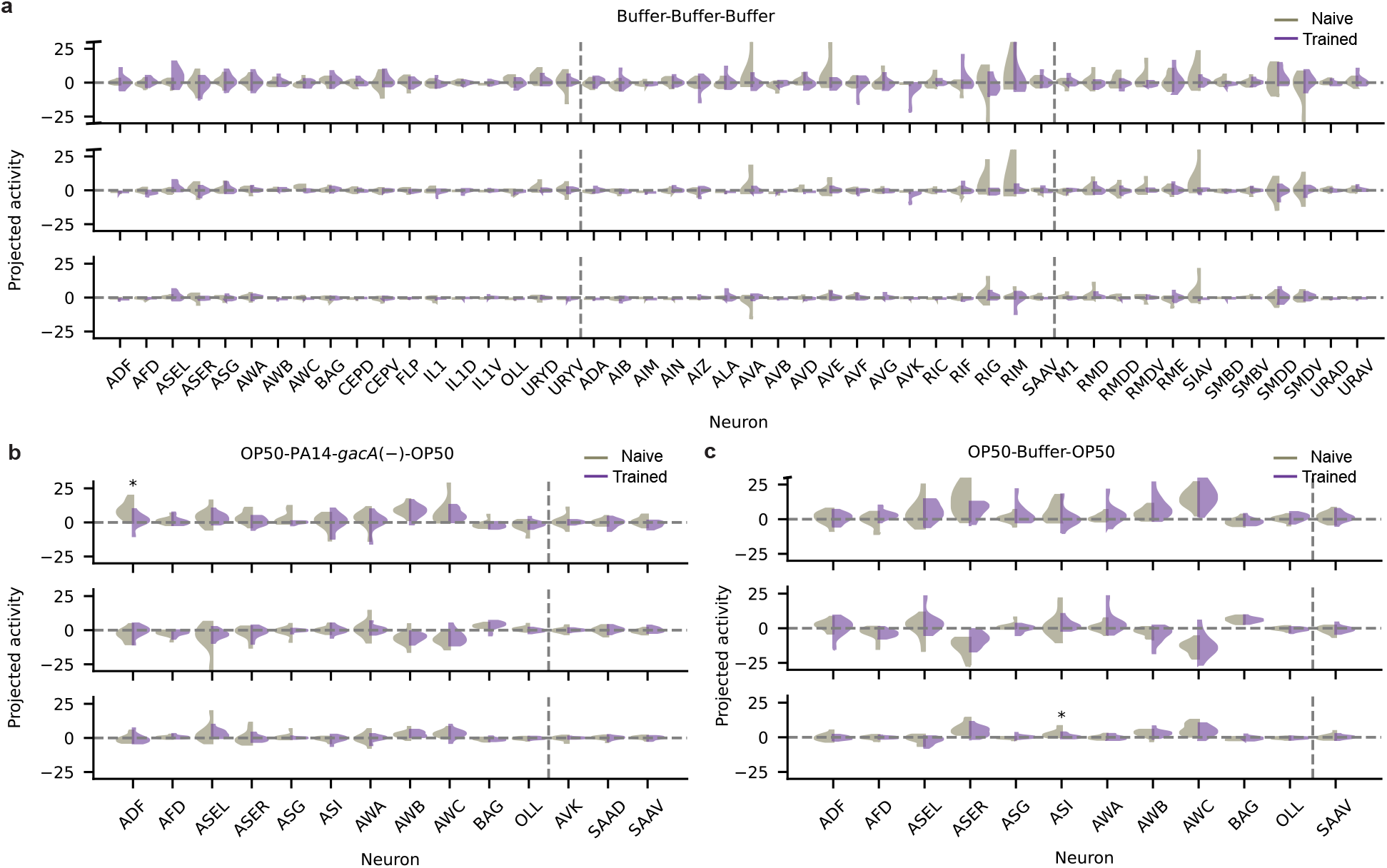
Violin-plots of the projections of neural responses from Buffer-Buffer-Buffer stimulation pattern (a), OP50-PA14-*gacA(-)*-OP50 stimulation pattern (b), and OP50-Buffer-OP50 stimulation pattern (c) to the 3 Temporal Components shown in (Figure 2). Each violin-plot represents the kernel density estimate for the distribution of the projections. Asterisks indicate neurons that were significantly different between naive (gray) and trained (purple) worms (Methods).

**Figure S9.**
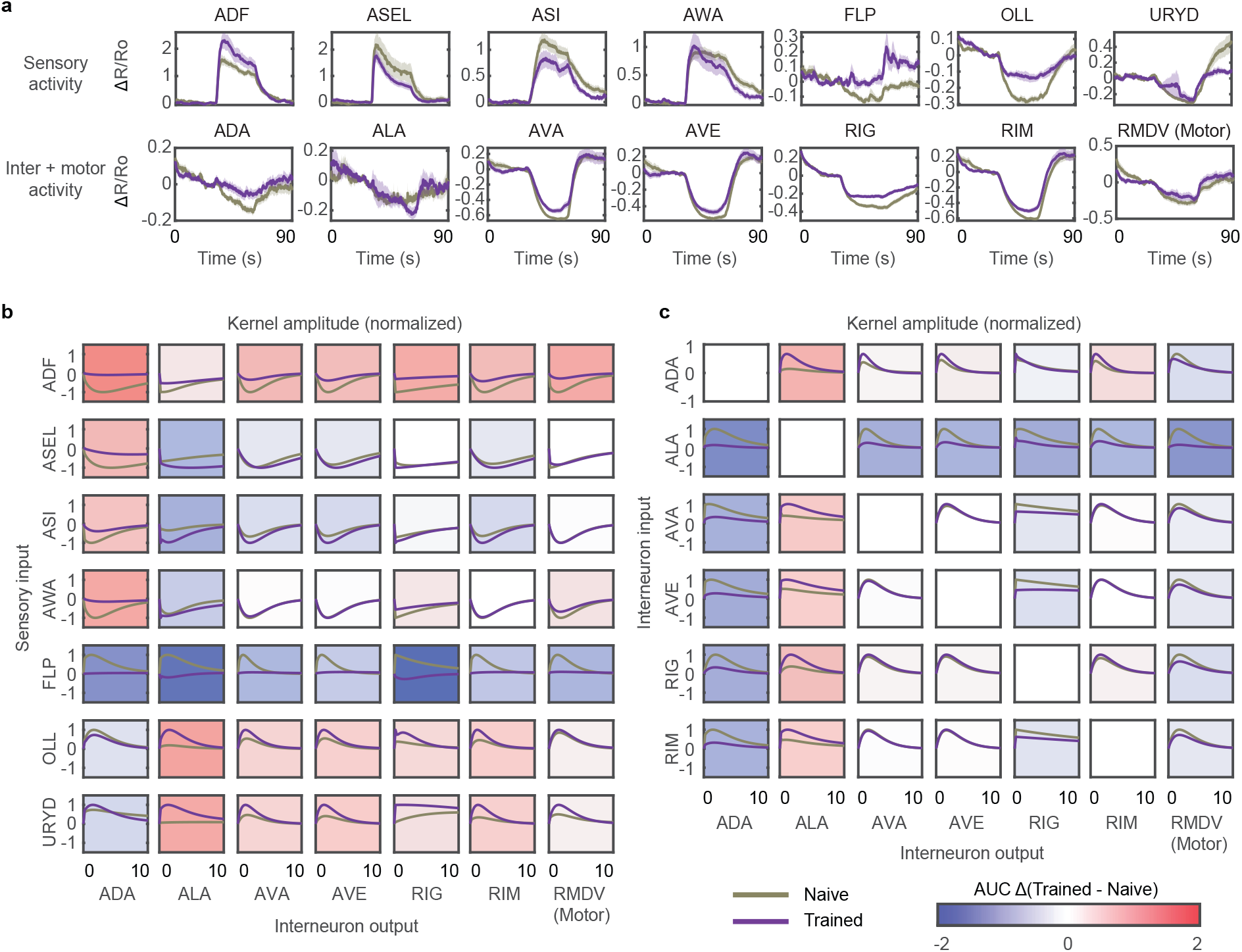
Changes in input-output relationship between the activities of sensory neurons, interneurons, and motor neurons are context-gated and reduced in the detection task. (a) Average activity and SEM (shaded region) of the neurons show in Figure 3a, but for the detection task (Buffer-PA14-Buffer stimulation pattern). (b) Double exponential kernels fitted between sensory (rows) and inter/motor neuron (column) activities show reduced divergence between naive and trained under the detection task, compared to extensive divergence seen in the discrimination task in Figure 3b. Kernels shown are the average across 2,000 bootstraps and normalized by maximum absolute value of amplitude across naive and trained kernels for visualization (normalization preserves relative scaling of naive and trained). Background color indicates normalized change in average kernel amplitude (AUC Δ) between naive and trained. (c) Kernels fitted between interneurons for detection task also demonstrate reduced divergence between naive and trained, compared to the divergence seen in the discrimination task in Figure 3c.

**Figure S10.**
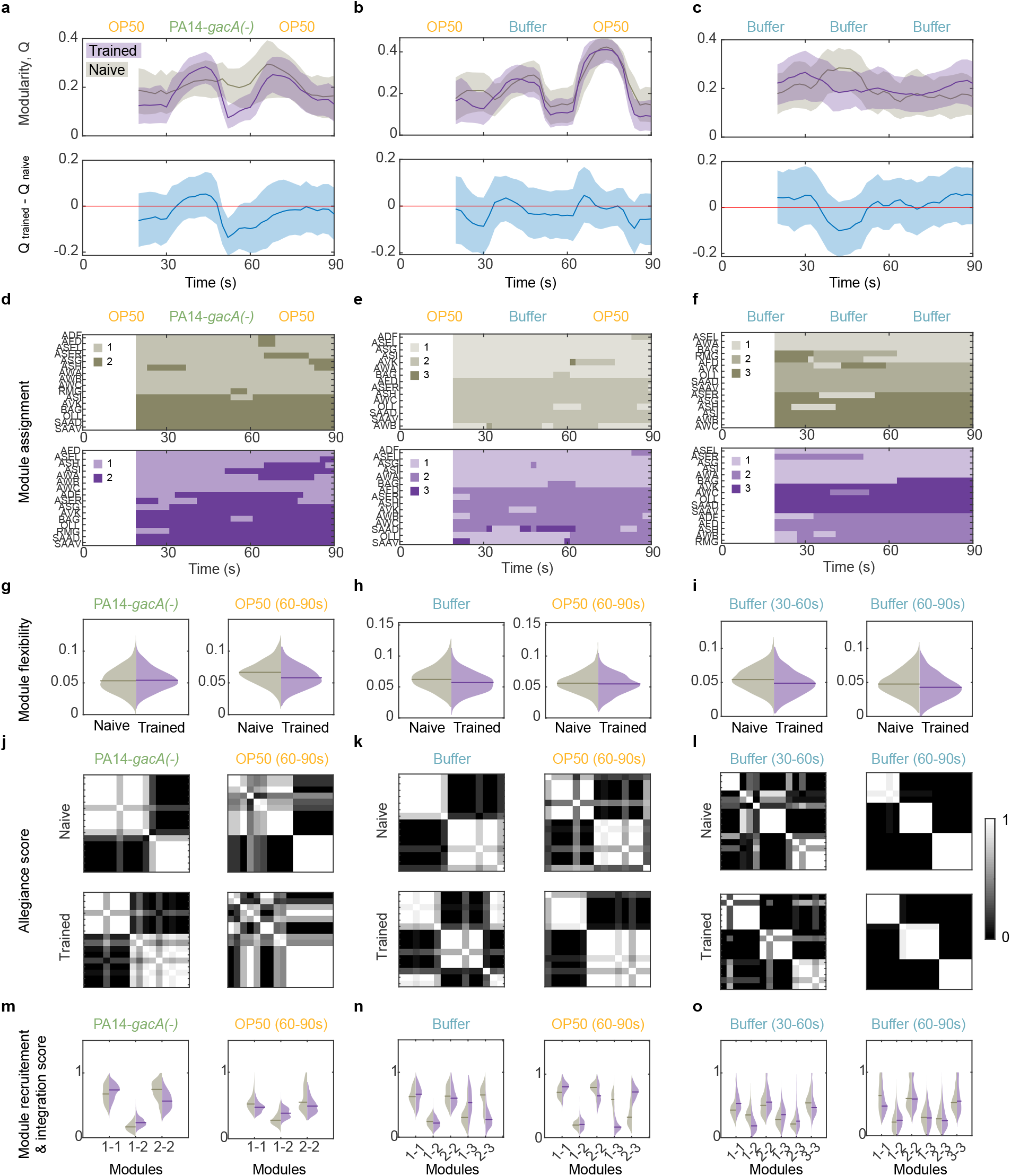
Changes in network organization of sensory–inter group I neurons are minimal outside of the bacterial odorant discrimination task. Analysis of network modularity (a–c), module assignments (d–f), network flexibility (g–i), allegiance matrices (j–l), and recruitment/integration (m–o) for additional stimulation patterns (column 1: OP50–PA14-*gacA(-)*–OP50; column 2: OP50–Buffer–OP50; column 3: Buffer–Buffer–Buffer). (a–c) Difference in naive and trained modularity is minimal across all three stimulation patterns, with only OP50–PA14-*gacA(-)*–OP50 showing a minor change at 50 s (a). (d–f) Changes in module recruitment patterns are minimal across all stimulation patterns. Naive and trained recruit the same number of modules in response to each pattern. Some reorganization is observed in the OP50–Buffer–OP50 context (e), with OLL and SAAV switching membership from module 2 in naive to module 1 during Buffer (30–60 s) in trained. (g–i) Network flexibility is stable across naive and trained (p = 0.44, 0.41, 0.40, 0.49, 0.42, 0.42 from left to right). Violin plots show the distribution of bootstrapped values (n = 1,000 bootstraps) and the median (horizontal line) for naive (gray) and trained (purple). Significance values were obtained from bootstrapped confidence intervals. (j–l) Stimulus-specific allegiance matrices of naive and trained module assignments. (m–o) Recruitment and integration between modules remain stable after learning across all three stimulation patterns, with only the OP50–Buffer–OP50 context (n) showing changes in integration between modules 1 and 2 and module 3 due to OLL and SAAV changing membership from module 2 in naive to module 3 in trained. Violin plots show the distribution of bootstrapped values (n = 1,000) and the median (horizontal line). Significance values were obtained using bootstrapped confidence intervals (n = 1,000) of naive and trained functional connectivity. Values are listed from left to right within each panel: (m, left) p = 0.42, 0.26, 0.22; (m, right) p = 0.34, 0.22, 0.33; (n, left) p = 0.46, 0.42, 0.46, 0.22, 0.09; (n, right) p = 0.31, 0.45, 0.29, 0.04, 0.09; (o, left) p = 0.27, 0.14, 0.14, 0.35, 0.40, 0.40; (o, right) p = 0.34, 0.46, 0.49, 0.49, 0.46, 0.47.

**Figure S11.**
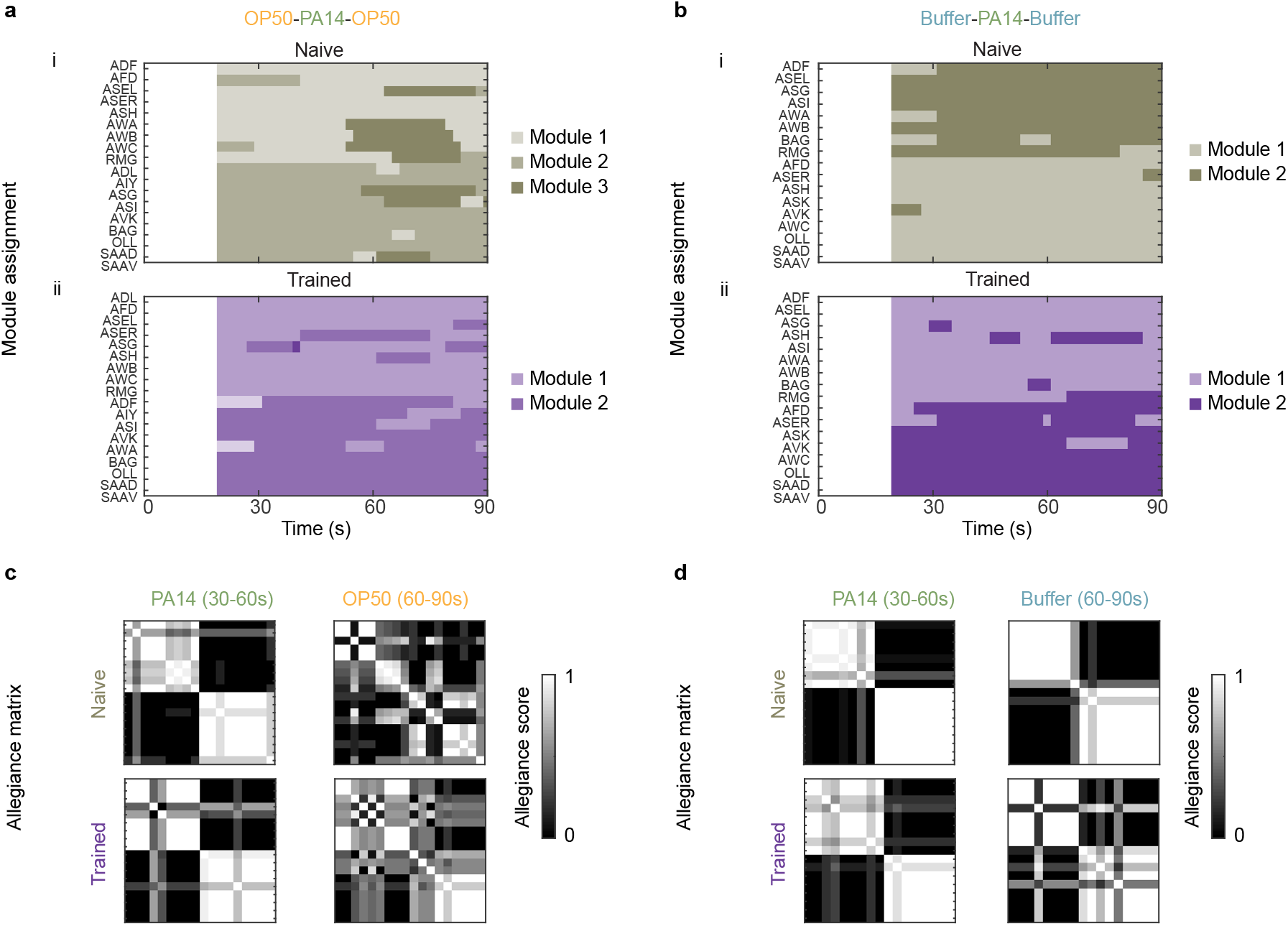
Module assignments and allegiance matrices for sensory-inter group I neurons during discrimination and detection tasks. (a) During discrimination task (OP50-PA14-OP50 stimulation pattern), naive activity organizes into two main modules for PA14 onset transitions (i, 30-60s) but three main modules for PA14 removal transitions (60-90s). After learning (ii), modules recruited during the discrimination task reorganize and PA14 onset (30-60s) and removal (60-90s) transitions recruit consistent sets of modules. Module assignments shown are a representative consensus assignment across n = 1,000 bootstraps. (b) During detection task (Buffer-PA14-Buffer stimulation pattern), modules recruited by PA14 onset (30-60s) and removal (60-90s) transitions are consistent in both naive (i) and trained (ii) and reorganization during learning is minimal (only ASH changes from primarily associating with module 2 in naive to module 1 in trained). (c, d) Stimulus specific allegiance matrices for naive and trained module assignments shown in (a, b) show pairwise probabilities of module co-assignment between neurons (allegiance score) for specific stimulus periods. For the discrimination task (c), recruitment patterns are distinct across stimulus periods in naive (c, top row), but stable across stimulus periods in trained (c, bottom row). For the detection task (d), recruitment patterns are relatively stable across stimulus periods and training, with slight destabilization of trained recruitment patterns in response to PA14 removal (d, lower right). Ordering of allegiance matrix rows and columns is consistent with corresponding module assignment heatmap in a,b.

**Figure S12.**
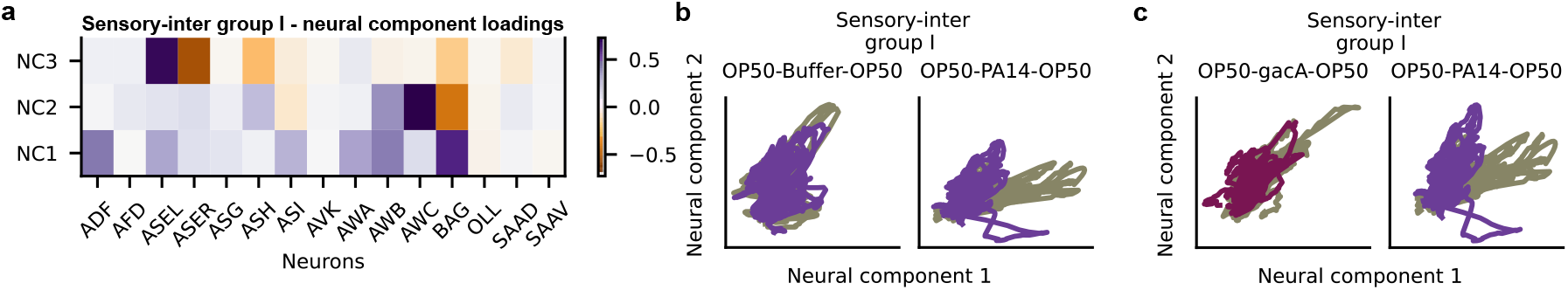
Rotation of the neural state manifolds is stimulus and training condition specific. (a) Loadings for the neural components 1-3 for the sensory-inter group I showing the principal axes of neural co-activations and their directions. Component 1 heavily weighs a subset of the amphid sensory neurons (ADF, ASEL, ASI, AWA, AWB, BAG), component 2 heavily weighs AWB, AWC and their reciprocal relation with BAG. (b,c) Neural activities of sensory-inter group I neurons in response to OP50-Buffer-OP50 stimulation pattern (b) or OP50-PA14-*gacA(-)*-OP50 (referred to as OP50-gacA-OP50) stimulation pattern (c) from naive (gray) and trained (purple, PA14-trained in **b**, and magenta, PA14-*gacA(-)* trained in **c**) animals projected to the Neural Components 1 and 2. For each panel, the data was projected using the loadings of the subset of neurons in the neural components that were common to the pair of datasets.

**Figure S13.**
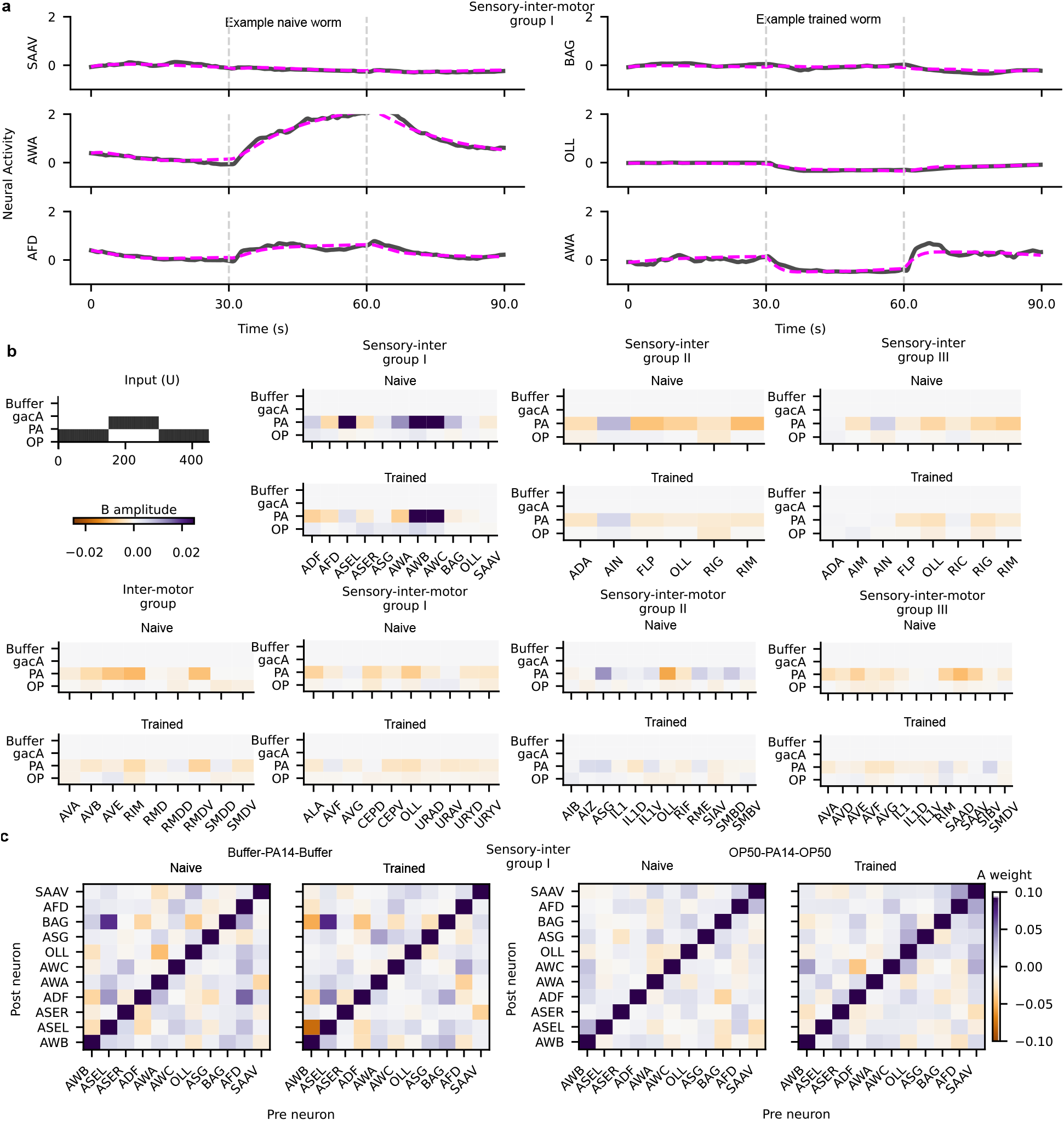
Fitting a Linear Dynamical System (LDS) model to neural activity data for naive and trained worms expressing GCaMP6 in different neuron groups, exposed to different odorant stimulation patterns. (Methods) (a) Example fits (magenta) to the neural data (black) of naive (left) and trained (right) worms for a few sample neurons. (b) *B* (input scaling) matrix averaged across worms belonging to the same neuron group and training condition and exposed to OP50-PA14-OP50 stimulation pattern (step-function of input shown top, left). Colors indicate the amplitude and sign for B showing both consistent responses and changes in input-responsiveness of neurons over learning. (c) Example *A* (cross-interactions) matrix averaged across naive and trained worms expressing GCaMP6 in sensory-inter group I neurons for Buffer-PA14-Buffer (left) and OP50-PA14-OP50 stimulation patterns (right).

**Figure S14.**
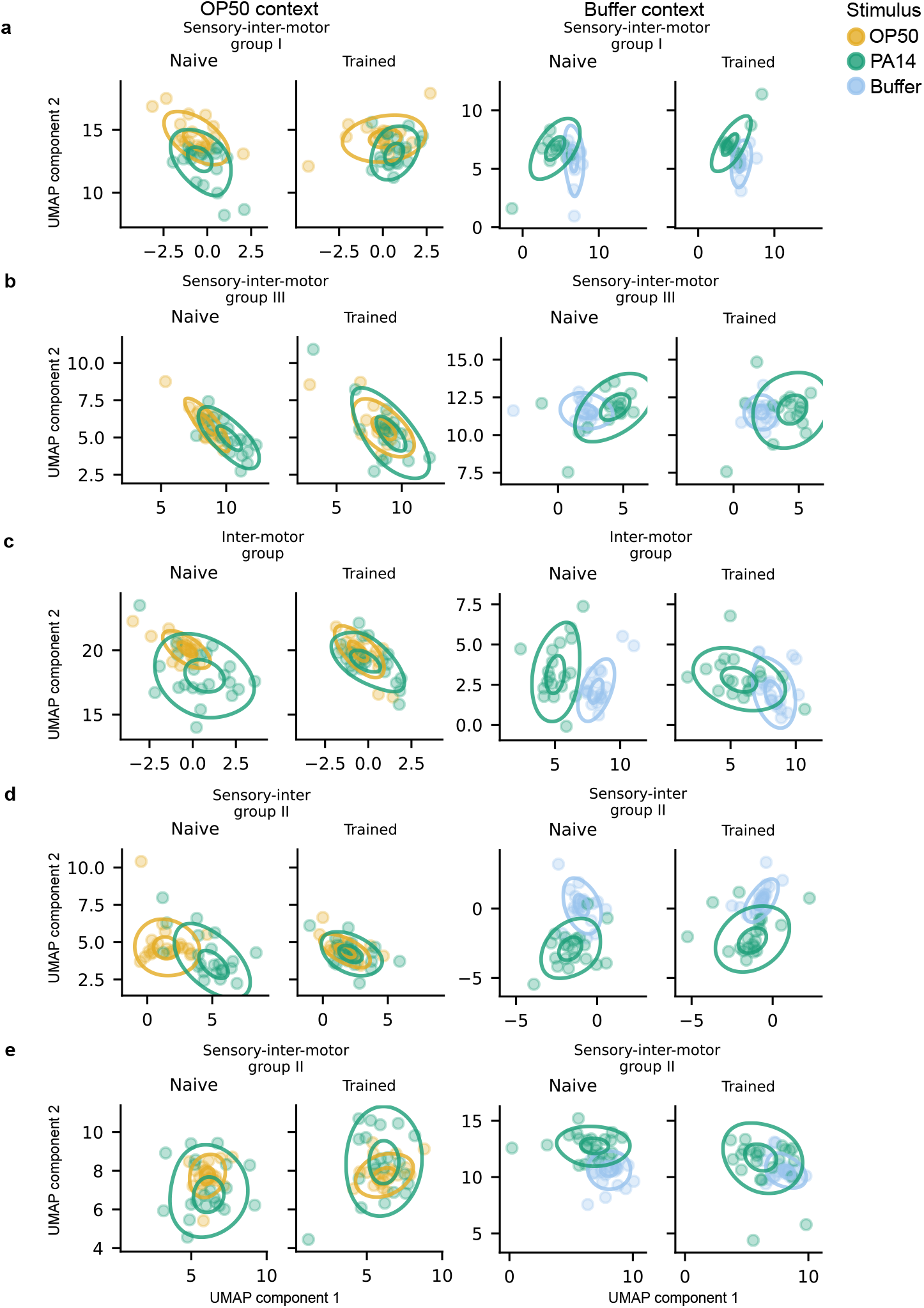
Shifts in fixed-points determined by the Linear Dynamical System model over learning for worms expressing GCaMP6 in different neuron groups (Methods) (a-e) UMAPs for different neuron groups under OP50 (left) and Buffer (right) contexts for naive and trained animals (Methods, same as Figure 5b-e). Each point is the fixed point for an individual worm when exposed to a given odor stimulation (OP50 or PA14 or Buffer) indicated by the color. Outer and inner ellipses are contours of kernel density estimates that enclose 90% and 50% of the distribution of fixed points respectively. Centroids of fixed-point distributions of the trained worms converge with the fixed point distributions of naive worms in a context dependent manner.

**Supplementary Table 1.** Geometric analysis of the low-dimensional neural state.

| Neuron group | stimulus | *R* | *q*<sup>*R*</sup> | *C* | *q*<sup>*C*</sup> |
|---|---|---|---|---|---|
| inter-motor group | Buffer-PA14-Buffer | 9.10 | 1.00 | 1.49 | 0.43 |
| **inter-motor group** | OP50-PA14-OP50 | 15.26 | 0.23 | 0.33 | **0.02** |
| sensory-inter group I | Buffer-PA14-Buffer | 2.96 | 1.00 | 0.85 | 0.36 |
| **sensory-inter group I** | OP50-PA14-OP50 | 44.50 | **0.00** | 1.76 | 0.55 |
| sensory-inter group II | Buffer-PA14-Buffer | 4.50 | 1.00 | 0.99 | 0.43 |
| **sensory-inter group II** | OP50-PA14-OP50 | 13.45 | **0.00** | 0.02 | **0.00** |
| sensory-inter group III | Buffer-PA14-Buffer | 5.11 | 1.00 | 1.06 | 0.41 |
| **sensory-inter group III** | OP50-PA14-OP50 | 150.33 | **0.00** | 0.17 | **0.00** |
| sensory-inter-motor group I | Buffer-PA14-Buffer | 0.95 | 1.00 | 0.65 | 0.88 |
| sensory-inter-motor group I | OP50-PA14-OP50 | 82.52 | 0.11 | 1.00 | 0.41 |
| sensory-inter-motor group II | Buffer-PA14-Buffer | 31.02 | 0.13 | 0.28 | 0.07 |
| sensory-inter-motor group II | OP50-PA14-OP50 | 151.04 | 0.13 | 0.44 | 0.36 |
| sensory-inter-motor group III | Buffer-PA14-Buffer | 1.69 | 1.00 | 0.86 | 0.41 |
| **sensory-inter-motor group III** | OP50-PA14-OP50 | 115.65 | **0.02** | 0.18 | 0.15 |

## References

1. Kandel, E. R., Dudai, Y. & Mayford, M. R. The Molecular and Systems Biology of Memory. Cell 157, 163–186. issn: 0092-8674. 10.1016/j.cell.2014.03.001 (Mar. 2014).

2. Josselyn, S. A. & Frankland, P. W. Memory Allocation: Mechanisms and Function. Annual Review of Neuroscience 41, 389–413. issn: 1545-4126. 10.1146/annurev-neuro-080317-061956 (July 2018).

3. Zhang, Y., Iino, Y. & Schafer, W. R. Behavioral plasticity. GENETICS 228 (ed Sengupta, P.) issn: 1943-2631. 10.1093/genetics/iyae105 (Aug. 2024).

4. Cognigni, P., Felsenberg, J. & Waddell, S. Do the right thing: neural network mechanisms of memory formation, expression and update in Drosophila. Current Opinion in Neurobiology 49, 51–58. issn: 0959-4388. 10.1016/j.conb.2017.12.002 (Apr. 2018).

5. Banerjee, A. et al. Value-guided remapping of sensory cortex by lateral orbitofrontal cortex. Nature 585, 245–250. issn: 1476-4687. 10.1038/s41586-020-2704-z (Sept. 2020).

6. Harvey, C. D., Coen, P. & Tank, D. W. Choice-specific sequences in parietal cortex during a virtual-navigation decision task. Nature 484, 62–68. issn: 1476-4687. 10.1038/nature10918 (Mar. 2012).

7. Reijmers, L. G., Perkins, B. L., Matsuo, N. & Mayford, M. Localization of a Stable Neural Correlate of Associative Memory. Science 317, 1230–1233. issn: 1095-9203. 10.1126/science.1143839 (Aug. 2007).

8. Han, J.-H. et al. Neuronal Competition and Selection During Memory Formation. Science 316, 457–460. issn: 1095-9203. 10.1126/science.1139438 (Apr. 2007).

9. Findling, C. et al. Brain-wide representations of prior information in mouse decision-making. Nature 645, 192–200. issn: 1476-4687. 10.1038/s41586-025-09226-1 (Sept. 2025).

10. Meshulam, L. et al. A brain-wide map of neural activity during complex behaviour. Nature 645, 177–191. issn: 1476-4687. 10.1038/s41586-025-09235-0 (Sept. 2025).

11. Ha, H.-i. et al. Functional Organization of a Neural Network for Aversive Olfactory Learning in Caenorhabditis elegans. Neuron 68, 1173–1186. issn: 0896-6273. 10.1016/j.neuron.2010.11.025 (Dec. 2010).

12. Zhang, Y., Lu, H. & Bargmann, C. I. Pathogenic bacteria induce aversive olfactory learning in Caenorhabditis elegans. Nature 438, 179–184. issn: 1476-4687. 10.1038/nature04216 (Nov. 2005).

13. Tan, M.-W., Mahajan-Miklos, S. & Ausubel, F. M. Killing of Caenorhabditis elegans by Pseudomonas aeruginosa used to model mammalian bacterial pathogenesis. Proceedings of the National Academy of Sciences 96, 715–720. issn: 1091-6490. 10.1073/pnas.96.2.715 (Jan. 1999).

14. Garcia, J., Kimeldorf, D. J. & Koellino, R. Conditioned Aversion to Saccharin Resulting from Exposure to Gamma Radiation. Science 122, 157–158. http://www.jstor.org/stable/1752118 (July 1955).

15. Garcia, J. & Koelling, R. A Comparison of Aversions Induced by X Rays, Toxins, and Drugs in the Rat. Radiation Research Supplement 7, 439–450. https://www.jstor.org/stable/3583736 (Sept. 1967).

16. Gershon, M. Plasticity in serotonin control mechanisms in the gut. Current Opinion in Pharmacology 3, 600–607. issn: 1471-4892. 10.1016/j.coph.2003.07.005 (Dec. 2003).

17. Luo, L., Gabel, C. V., Ha, H.-I., Zhang, Y. & Samuel, A. D. T. Olfactory Behavior of Swimming C. elegans Analyzed by Measuring Motile Responses to Temporal Variations of Odorants. Journal of Neurophysiology 99, 2617–2625. issn: 1522-1598. 10.1152/jn.00053.2008 (May 2008).

18. Pierce-Shimomura, J. T., Morse, T. M. & Lockery, S. R. The Fundamental Role of Pirouettes in Caenorhabditis elegans Chemotaxis. The Journal of Neuroscience 19, 9557–9569. issn: 1529-2401. 10.1523/JNEUROSCI.19-21-09557.1999 (Nov. 1999).

19. Iino, Y. & Yoshida, K. Parallel Use of Two Behavioral Mechanisms for Chemotaxis in Caenorhabditis elegans. The Journal of Neuroscience 29, 5370–5380. issn: 1529-2401. 10.1523/JNEUROSCI.3633-08.2009 (Apr. 2009).

20. Chen, T.-W. et al. Ultrasensitive fluorescent proteins for imaging neuronal activity. Nature 499, 295–300. issn: 1476-4687. 10.1038/nature12354 (July 2013).

21. Shaner, N. C. et al. Improved monomeric red, orange and yellow fluorescent proteins derived from Discosoma sp. red fluorescent protein. Nature Biotechnology 22, 1567–1572. issn: 1546-1696. 10.1038/nbt1037 (Nov. 2004).

22. Altun, Z. et al. WormAtlas. http://www.wormatlas.org (2002x-2024).

23. Bargmann, C. I., Hartwieg, E. & Horvitz, H. R. Odorant-selective genes and neurons mediate olfaction in C. elegans. Cell 74, 515–527. issn: 0092-8674. 10.1016/0092-8674(93)80053-h (Aug. 1993).

24. Gordus, A., Pokala, N., Levy, S., Flavell, S. W. & Bargmann, C. I. Feedback from Network States Generates Variability in a Probabilistic Olfactory Circuit. Cell 161, 215–227. issn: 0092-8674. 10.1016/j.cell.2015.02.018 (Apr. 2015).

25. Gray, J. M., Hill, J. J. & Bargmann, C. I. A circuit for navigation in Caenorhabditis elegans. Proceedings of the National Academy of Sciences 102, 3184–3191. issn: 1091-6490. 10.1073/pnas.0409009101 (Feb. 2005).

26. Hendricks, M., Ha, H., Maffey, N. & Zhang, Y. Compartmentalized calcium dynamics in a C. elegans interneuron encode head movement. Nature 487, 99–103. issn: 1476-4687. 10.1038/nature11081 (May 2012).

27. Brissette, B. et al. Chemosensory detection of polyamine metabolites guides C. elegans to nutritive microbes. Science Advances 10. issn: 2375-2548. 10.1126/sciadv.adj4387 (Mar. 2024).

28. Wu, T. et al. Pheromones Modulate Learning by Regulating the Balanced Signals of Two Insulin-like Peptides. Neuron 104, 1095–1109.e5. issn: 0896-6273. 10.1016/j.neuron.2019.09.006 (Dec. 2019).

29. White, J. G., Southgate, E., J.N., T. & Brenner, S. Philosophical Transactions of the Royal Society of London. B, Biological Sciences 314, 1–340. issn: 2054-0280. 10.1098/rstb.1986.0056 (Nov. 1986).

30. Kaplan, H. S., Salazar Thula, O., Khoss, N. & Zimmer, M. Nested Neuronal Dynamics Orchestrate a Behavioral Hierarchy across Timescales. Neuron 105, 562–576.e9. issn: 0896-6273. 10.1016/j.neuron.2019.10.037 (Feb. 2020).

31. Shen, Y. et al. An extrasynaptic GABAergic signal modulates a pattern of forward movement in Caenorhabditis elegans. eLife 5. issn: 2050-084X. 10.7554/eLife.14197 (May 2016).

32. Kato, S. et al. Global Brain Dynamics Embed the Motor Command Sequence of Caenorhabditis elegans. Cell 163, 656–669. issn: 0092-8674. 10.1016/j.cell.2015.09.034 (Oct. 2015).

33. Atanas, A. A. et al. Brain-wide representations of behavior spanning multiple timescales and states in C. elegans. Cell 186, 4134–4151.e31. issn: 0092-8674. 10.1016/j.cell.2023.07.035 (Sept. 2023).

34. Brezovec, B. E. et al. Mapping the neural dynamics of locomotion across the Drosophila brain. Current Biology 34, 710–726.e4. issn: 0960-9822. 10.1016/j.cub.2023.12.063 (Feb. 2024).

35. Zhang, X. & Zhang, Y. DBL-1, a TGF-, is essential for Caenorhabditis elegans aversive olfactory learning. Proceedings of the National Academy of Sciences 109, 17081–17086. issn: 1091-6490. 10.1073/pnas.1205982109 (Sept. 2012).

36. Rahme, L. G. et al. Use of model plant hosts to identify Pseudomonas aeruginosa virulence factors. Proceedings of the National Academy of Sciences 94, 13245–13250. issn: 1091-6490. 10.1073/pnas.94.24.13245 (Nov. 1997).

37. Newman, M. E. J. Modularity and community structure in networks. Proceedings of the National Academy of Sciences 103. Original modularity paper, 8577–8582. issn: 0027-8424. eprint: physics/0602124 (2006).

38. Bassett, D. S., Yang, M., Wymbs, N. F. & Grafton, S. T. Learning-induced autonomy of sensorimotor systems. Nature Neuroscience 18, 744–751. issn: 1097-6256 (2015).

39. Bassett, D. S. et al. Dynamic reconfiguration of human brain networks during learning. Proceedings of the National Academy of Sciences 108, 7641–7646. issn: 0027-8424. eprint: 1010.3775 (2011).

40. Chung, S. & Abbott, L. F. Neural population geometry: An approach for understanding biological and artificial neural networks. Current opinion in neurobiology 70, 137–144 (2021).

41. Vyas, S., Golub, M. D., Sussillo, D. & Shenoy, K. V. Computation through neural population dynamics. Annual review of neuroscience 43, 249–275 (2020).

42. Kao, T.-C. & Hennequin, G. Neuroscience out of control: control-theoretic perspectives on neural circuit dynamics. Current Opinion in Neurobiology 58, 122–129. issn: 0959-4388. 10.1016/j.conb.2019.09.001 (Oct. 2019).

43. Warden, M. R. & Miller, E. K. Task-Dependent Changes in Short-Term Memory in the Prefrontal Cortex. The Journal of Neuroscience 30, 15801–15810. issn: 1529-2401. 10.1523/JNEUROSCI.1569-10.2010 (Nov. 2010).

44. Rigotti, M. et al. The importance of mixed selectivity in complex cognitive tasks. Nature 497, 585–590. issn: 1476-4687. 10.1038/nature12160 (May 2013).

45. Dang, W., Jaffe, R. J.Qi, X.-L. & Constantinidis, C. Emergence of Non-Linear Mixed Selectivity in Prefrontal Cortex after Training. The Journal of Neuroscience, JN–RM–2814–20. issn: 1529-2401. 10.1523/JNEUROSCI.2814-20.2021 (July 2021).

46. Faulkner, A. D. et al. Context flexibly modulates cue representations in visual cortex. Nature Communications 16. issn: 2041-1723. 10.1038/s41467-025-61314-y (July 2025).

47. Harris, K. D. & Thiele, A. Cortical state and attention. Nature Reviews Neuroscience 12, 509–523. issn: 1471-0048. 10.1038/nrn3084 (Aug. 2011).

48. Gilbert, C. D. & Li, W. Top-down influences on visual processing. Nature Reviews Neuroscience 14, 350–363. issn: 1471-0048. 10.1038/nrn3476 (Apr. 2013).

49. Schneider, D. M. & Mooney, R. How Movement Modulates Hearing. Annual Review of Neuroscience 41, 553–572. issn: 1545-4126. 10.1146/annurev-neuro-072116-031215 (July 2018).

50. McCloskey, M. & Cohen, N. J. in Psychology of learning and motivation 109–165 (Elsevier, 1989).

51. Masse, N. Y., Grant, G. D. & Freedman, D. J. Alleviating catastrophic forgetting using context-dependent gating and synaptic stabilization. Proceedings of the National Academy of Sciences 115, E10467–E10475 (2018).

52. Brenner, S. The genetics of Caenorhabditis elegans. Genetics 77, 71–94 (1974).

53. Brockie, P. J., Mellem, J. E., Hills, T., Madsen, D. M. & Maricq, A. V. The C. elegans glutamate receptor subunit NMR-1 is required for slow NMDA-activated currents that regulate reversal frequency during locomotion. Neuron 31, 617–630 (2001).

54. Winnier, A. R. et al. UNC-4/UNC-37-dependent repression of motor neuron-specific genes controls synaptic choice in Caenorhabditis elegans. Genes & development 13, 2774–2786 (1999).

55. Macosko, E. Z. et al. A hub-and-spoke circuit drives pheromone attraction and social behaviour in C. elegans. Nature 458, 1171–1175 (2009).

56. Altun, Z. F., Chen, B.Wang, Z.-W. & Hall, D. H. High resolution map of Caenorhabditis elegans gap junction proteins. Developmental dynamics: an official publication of the American Association of Anatomists 238, 1936–1950 (2009).

57. Kage, E. et al. MBR-1, a novel helix-turn-helix transcription factor, is required for pruning excessive neurites in Caenorhabditis elegans. Current Biology 15, 1554–1559 (2005).

58. Brockie, P. J., Madsen, D. M., Zheng, Y., Mellem, J. & Maricq, A. V. Differential expression of glutamate receptor subunits in the nervous system of Caenorhabditis elegans and their regulation by the homeodomain protein UNC-42. Journal of Neuroscience 21, 1510–1522 (2001).

59. Majeed, M. et al. Toolkits for detailed and high-throughput interrogation of synapses in C. elegans. Elife 12, RP91775 (2024).

60. Berghoff, E. G. et al. The Prop1-like homeobox gene unc-42 specifies the identity of synaptically connected neurons. Elife 10, e64903 (2021).

61. Troemel, E. R., Chou, J. H., Dwyer, N. D., Colbert, H. A. & Bargmann, C. I. Divergent seven transmembrane receptors are candidate chemosensory receptors in C. elegans. Cell 83, 207–218 (1995).

62. Chou, J. H., Bargmann, C. I. & Sengupta, P. The Caenorhabditis elegans odr-2 gene encodes a novel Ly-6-related protein required for olfaction. Genetics 157, 211–224 (2001).

63. Mello, C. C., Kramer, J. M., Stinchcomb, D. & Ambros, V. Efficient gene transfer in C. elegans: extrachromosomal maintenance and integration of transforming sequences. The EMBO journal 10, 3959–3970 (1991).

64. Lee, H. J. et al. Automated cell annotation in multi-cell images using an improved CRF_ID algorithm. eLife 12, RP89050 (2025).

65. Reddy, K. C., Andersen, E. C., Kruglyak, L. & Kim, D. H. A polymorphism in npr-1 is a behavioral determinant of pathogen susceptibility in C. elegans. Science 323, 382–384 (2009).

66. Chronis, N., Zimmer, M. & Bargmann, C. I. Microfluidics for in vivo imaging of neuronal and behavioral activity in Caenorhabditis elegans. Nature Methods 4, 727–731. issn: 1548-7105. 10.1038/nmeth1075 (Aug. 2007).

67. Thevenaz, P., Ruttimann, U. E. & Unser, M. A pyramid approach to subpixel registration based on intensity. IEEE transactions on image processing 7, 27–41 (1998).

68. Suzuki, H. et al. Functional asymmetry in Caenorhabditis elegans taste neurons and its computational role in chemotaxis. Nature 454, 114–117. issn: 1476-4687. 10.1038/nature06927 (July 2008).

69. Mucha, P. J., Richardson, T., Macon, K., Porter, M. A. & Onnela, J.-P. Community Structure in Time-Dependent, Multiscale, and Multiplex Networks. Science 328, 876–878. issn: 0036-8075. eprint: 0911.1824 (2010).

70. Blondel, V. D., Guillaume, J.-L., Lambiotte, R. & Lefebvre, E. Fast unfolding of communities in large networks. Journal of Statistical Mechanics: Theory and Experiment 2008, P10008. eprint: 0803.0476 (2008).

71. Jeub, L. G. S., Bazzi, M., Jutla, I. S. & Mucha, P. J. GenLouvain: A generalized Louvain method for community detection implemented in MATLAB https://github.com/GenLouvain/GenLouvain (2025).

72. Gómez, S., Jensen, P. & Arenas, A. Analysis of community structure in networks of correlated data. Physical Review E 80, 016114. issn: 1539-3755. eprint: 0812.3030 (2009).

73. Bassett, D. S. et al. Robust detection of dynamic community structure in networks. Chaos: An Interdisciplinary Journal of Nonlinear Science 23, 013142. issn: 1054-1500. eprint: 1206.4358 (2013).

74. Moza, S. & Zhang, Y. CeDNe: A multi-scale computational framework for modeling structure-function relationships in the C. elegans nervous system. submitted to BioRxiv (2025).

